# A multiomics approach reveals RNA dynamics promote cellular sensitivity to DNA hypomethylation

**DOI:** 10.1101/2022.12.14.518457

**Authors:** Alex Y. Ge, Abolfazl Arab, Raymond Dai, Albertas Navickas, Lisa Fish, Kristle Garcia, Hosseinali Asgharian, Jackson Goudreau, Sean Lee, Kathryn Keenan, Melissa B. Pappalardi, Michael T. McCabe, Laralynne Przybyla, Hani Goodarzi, Luke A. Gilbert

**Affiliations:** School of Medicine, University of California, San Francisco, San Francisco, CA 94158, USA; Arc Institute, Palo Alto, CA 94304, USA; Department of Urology, University of California, San Francisco, San Francisco, CA 94158, USA; Helen Diller Family Comprehensive Cancer Center, University of California, San Francisco, San Francisco, CA 94158, USA; Department of Biochemistry and Biophysics, University of California, San Francisco, San Francisco, CA 94158, USA; Bakar Computational Health Sciences Institute, University of California, San Francisco, San Francisco, CA 94158, USA; Tumor Cell Targeting Research Unit, Research, GSK, Collegeville, PA 19426, USA; Laboratory for Genomics Research, San Francisco, CA 94158, USA

## Abstract

The search for new approaches in cancer therapy requires a mechanistic understanding of cancer vulnerabilities and anti-cancer drug mechanisms of action. Problematically, some effective therapeutics target cancer vulnerabilities that have poorly defined mechanisms of anti-cancer activity. One such drug is decitabine, a frontline therapeutic approved for the treatment of high-risk acute myeloid leukemia (AML). Decitabine is thought to kill cancer cells selectively via inhibition of DNA methyltransferase enzymes, but the genes and mechanisms involved remain unclear. Here, we apply an integrated multiomics and CRISPR functional genomics approach to identify genes and processes associated with response to decitabine in AML cells. Our integrated multiomics approach reveals RNA dynamics are key regulators of DNA hypomethylation induced cell death. Specifically, regulation of RNA decapping, splicing and RNA methylation emerge as important regulators of cellular response to decitabine.

## INTRODUCTION

Epigenetic dysregulation drives many of the hallmarks of cancer by enabling aberrant gene expression programs which underlie cancer cellular plasticity and tumor heterogeneity phenotypes that promote cancer initiation, progression, metastasis and drug resistance^1^. Indeed, one of the key findings of the genomics era in cancer biology has been that most cancer genomes are epigenetically abnormal and mutations in genes that regulate DNA methylation, such as *DNMT3A/B*, *TET1-3* and *IDH1/2*, are prevalent^2,3^. Together, these observations suggest that epigenetic dysregulation promotes cancer but may also represent a targetable vulnerability. As such, there has been substantial interest in the development of anti-cancer strategies which modulate cancer associated epigenetic programs and dependencies^4–6^. One such promising strategy which has shown success in the context of certain subtypes of acute myeloid leukemia (AML) is to inhibit the activity of key enzymes required for maintenance and regulation of DNA methylation by small molecule drugs, such as decitabine, resulting in global DNA hypomethylation. There is clear evidence of clinical benefit of decitabine treatment for AML patients who have cytogenetic abnormalities associated with unfavorable risk, *TP53* mutations or both (defined hereafter as high-risk AML patients)^7,8^. Unfortunately, despite this benefit, most AML patients eventually progress following decitabine treatment with a median overall survival of less than 1 year. Problematically, relatively little progress has been made on improving the clinical activity of DNA hypomethylating agents (HMA) such as decitabine in AML or other cancers in part because the molecular determinants of response to HMAs are unclear.

A recent clinical study of molecular determinants of response to decitabine in AML patients has suggested that mutations in *DNMT3A*, *IDH1/2* and *TET2* are not correlated with response to decitabine^8^. In the same study, it was noted that *TP53* mutations are also not correlated with poor clinical response to decitabine. These findings are unusual in two ways. First, it had previously been hypothesized that tumors with mutations that drive aberrant DNA methylation profiles may be more susceptible to HMAs. Secondly, *TP53* mutations are generally associated with drug resistance and poor prognosis in many cancers, so it is unexpected that *TP53* mutations in AML seem to not play a role in determining clinical outcomes following treatment with decitabine. This result suggests that decitabine’s anti-cancer activity in AML occurs through a *TP53* independent mechanism. Given the central role *TP53* plays in canonical apoptotic *BCL2* family protein dependent programmed cell death, at one level this study appears to contradict recent clinical trial results in AML which demonstrated superior clinical outcomes from the combination of HMAs and venetoclax, a *BCL2* inhibitor thought to drive programmed cell death in cancer cells^9^. One explanation that could account for both sets of clinical observations is that HMAs may drive cell death via an unknown *TP53* independent apoptotic pathway. A more robust understanding of decitabine’s mechanisms of anti-cancer activity in *TP53*-mutant tumors could enable innovative therapeutic strategies and a better understanding of patients who do and do not respond robustly to HMAs. An alternate hypothesis for how HMAs kill cancer cells arises from the observation that treatment with HMAs results in accumulation of non-canonical transcripts including inverted SINE elements, endogenous retroviral elements and cryptic transcription start sites encoded in long terminal repeats which collectively act to induce immune activation^10–14^. Lastly, it has also been suggested that HMAs induce cellular differentiation in AML which may contribute to therapeutic efficacy^15^.

To identify genes that modulate decitabine’s anti-cancer activity in high-risk AML in an unbiased manner, we performed genome-scale CRISPR genetic screens and integrated this data with multiomics measurements of decitabine response in AML cells. Our results recapitulate multiple known factors which modulate response to decitabine, including *DCK*, *SLC29A1*, *MCL1* and *BCL2*, indicating the utility and robustness of our approach for interrogating the biology of decitabine in AML^9,16–22^. Central to our study was the finding that epitranscriptomic RNA modification and RNA quality control pathways effectively modulate response to decitabine in AML cells. In short, we have identified unexpected regulatory connections between DNA methylation, RNA methylation and RNA quality control pathways, which may provide further insight into decitabine’s mechanism(s) of action.

## RESULTS

### A genome-scale CRISPRi screen in AML cells identifies genes modulating decitabine sensitivity and resistance

We set out to perform a genome-scale genetic screen using our previously described CRISPR interference (CRISPRi) functional genomics platform to identify genes that regulate cancer cell response to decitabine (Fig. 1a), a clinically approved HMA^23,24^. For this, we used the HL-60 cell line, which is an established model of AML. The cell line is *TP53*, *NRAS* and *MYC* mutant and captures the biology of high-risk AML and more generally of an aggressive human cancer. To begin, we generated an HL-60 CRISPRi cell-line model that stably expressed the dCas9-BFP-KRAB fusion protein. We validated that the resulting CRISPRi HL-60 cell line, hereafter referred to as HL-60i, is highly active for targeted gene knockdowns (Supplementary Fig. 1a).

**Figure 1.**
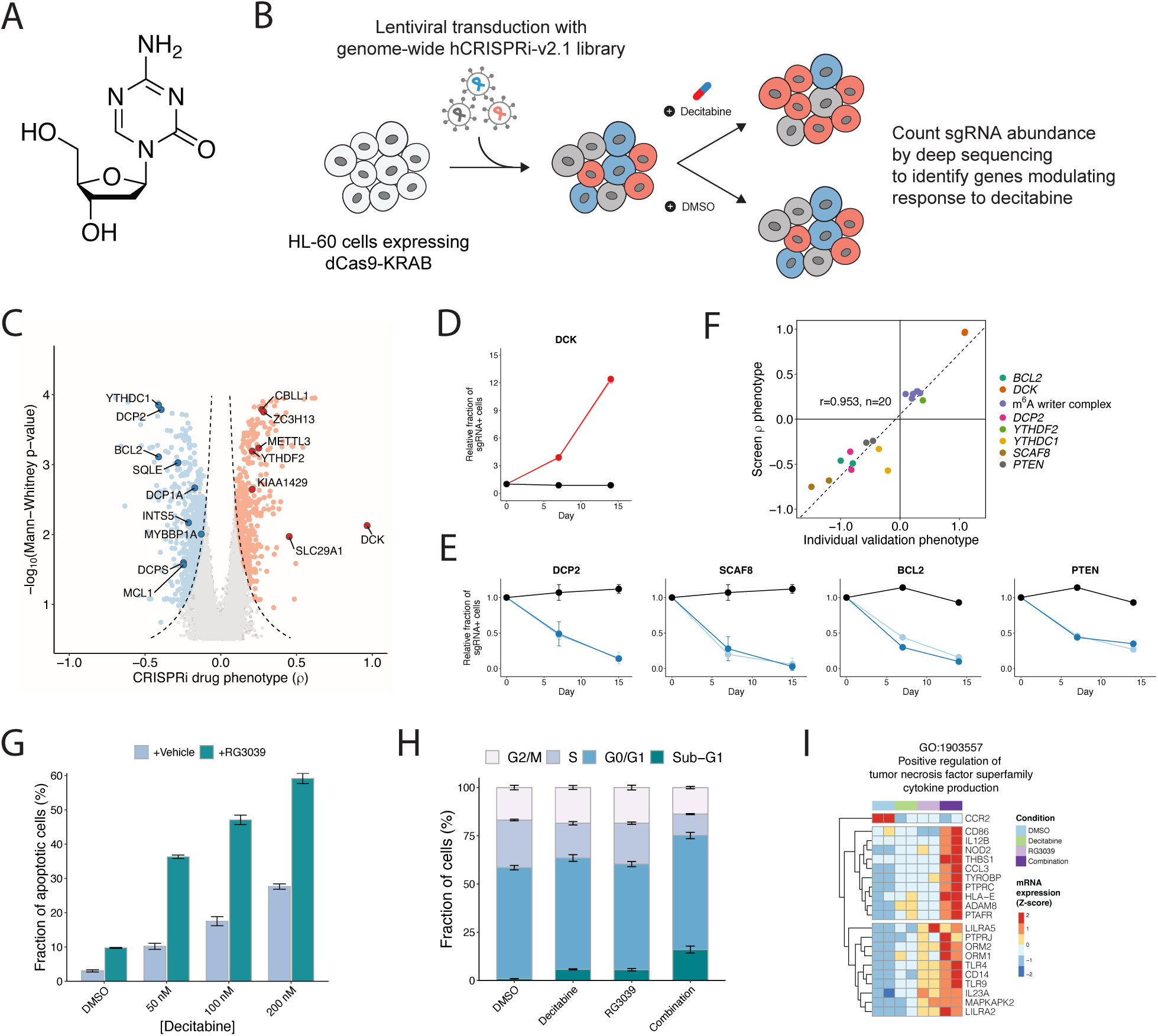
A genome-scale CRISPRi screen reveals gene knockdowns that confer sensitivity or resistance to 5-aza-2’-deoxycytidine (decitabine) (**a**) The chemical structure of decitabine. (**b**) Schematic of a genome-scale CRISPRi screen performed in HL-60 cells. (**c**) Volcano plot of gene-level rho (ρ) phenotypes and Mann-Whitney p-values. Negative rho values represent increased sensitivity to decitabine after knockdown, and positive rho values represent increased resistance. (**d-e**) Validation of top screen hits. HL-60i cells were transduced with a control sgRNA (black) or an active sgRNA (red or blue) and treated with DMSO or decitabine, and the proportion of sgRNA+ cells in the decitabine condition relative to DMSO was observed over time. Data are shown as means ± SD, two sgRNAs per gene and two replicates per sgRNA. (**f**) Scatter plot showing the correlation between screen rho phenotype and validation phenotype (day 14-15 post-infection) for each validated sgRNA. (**g**) A cleaved caspase 3/7 assay shows the fraction of apoptotic HL-60 cells at day 5 following treatment with DMSO or decitabine ± RG3039. Data are shown as means ± SD for three replicates. (**h**) A cell cycle assay shows the fraction of HL-60 cells at different phases of the cell cycle at day 5 following treatment with DMSO or decitabine ± RG3039. Data are shown as means ± SD for three replicates. (**i**) Normalized counts for genes in GO:1903557 (positive regulation of tumor necrosis factor superfamily cytokine production) upregulated upon decitabine and RG3039 treatment.

Decitabine (5-aza-2’-deoxycytidine) is a pro-drug that is converted intracellularly into 5-aza-2’-deoxycytidine monophosphate^17,19,22^. This nucleoside analogue is in turn incorporated into DNA during replication, where it is thought to irreversibly and covalently trap and inhibit DNA methyltransferases *DNMT1*/*DNMT3A*/*DNMT3B* (Fig. 1a). Trapping of DNMTs renders them enzymatically inactive, resulting in global DNA hypomethylation and dysregulated gene expression. This broad reprogramming of the gene expression landscape results in cell cycle arrest or cell death through poorly characterized molecular mechanisms. At high doses, decitabine also causes DNA replicative stress and DNA damage. To further characterize decitabine’s activity in an AML cell model, we used publicly available data to analyze changes in genome-scale DNA methylation patterns in HL-60 cells treated for 120 hours with a low dose of decitabine (Supplementary Fig. 1b-d)^25^. As expected, we observed global hypomethylation of CpG dinucleotides and hypomethylation of differentially methylated regions (DMRs) following treatment with decitabine. This confirms the expected activity of decitabine, a non-specific DNMT inhibitor, in AML cells. As discussed above, there is a hypothesis raised by clinical results that perhaps decitabine induces *TP53* independent but *BCL2* family protein dependent apoptosis. To address this, we next assessed whether decitabine treatment induces caspase 3/7 dependent apoptosis in our HL-60 model. We observed a dose dependent increase in caspase 3/7 activation upon treatment with low concentrations of decitabine (Supplementary Fig. 1e). Together, our results indicate that decitabine induces *TP53*-independent apoptosis and DNA hypomethylation in a model of high-risk AML and further supports our notion that this model could provide insight into decitabine’s mechanism(s) of action.

For the genome-scale CRISPRi screen design and all subsequent experiments, we chose to treat cells with a clinically relevant low dose of decitabine (∼IC30; 100 nM)^26^. At this concentration, decitabine’s anti-cancer activity is thought to predominantly arise due to global DNA hypomethylation rather than via DNA replication stress^27,28^. The genome-scale pooled genetic screen was performed by transducing the cell line with a human genome-scale CRISPRi sgRNA library at a low multiplicity of infection such that a single sgRNA is expressed in most cells, and then cells were selected with puromycin to remove uninfected cells from the population (Fig. 1b). In addition to time-zero samples, we also collected samples after growing the library in the presence and absence of decitabine (in biological duplicates). Next-generation sequencing was used to quantify the relative abundance of cells expressing each sgRNA in each sample. We then used measurements across the entire library to calculate sgRNA- and gene-level phenotypic scores (Supplementary Fig. 2a). Results obtained from the replicate screens were highly correlated with high data quality in both the DMSO and decitabine experiments (Supplementary Fig. 2b-e). Analysis of our decitabine screen data revealed a large number of genes that modulate cellular response to decitabine (1293 genes with Mann-Whitney p-value < 0.05 and absolute value of rho score > 0.1) (Fig. 1c and Supplementary Table 1). These results may reflect the pleiotropic nature of DNA methylation biology.

Initial inspection of top hits from our decitabine CRISPRi screen in HL-60 cells recapitulated a number of genes whose knockdown is known to impact drug resistance and sensitivity (Fig. 1c). For example, the top resistance hit was *DCK*, which phosphorylates decitabine resulting in conversion of the pro-drug to the active drug^18,19^. Another top resistance hit was *SLC29A1*, which is a solute carrier protein required for decitabine entry into cells^18,19^. Lastly, *DCTD* is thought to play a role in the metabolism of decitabine and is revealed as a strong resistance hit as well^29^. We also observed that knockdown of *BCL2* and *MCL1* sensitizes HL-60i to decitabine, as expected from the clinical literature which suggests decitabine induces *BCL2* family protein mediated cell death^20,21^. The recapitulation of known positive control hits in our screens indicate the utility and robustness of our approach for interrogating the biology of decitabine in AML.

### RNA dynamics modulate response to DNA hypomethylation induced by decitabine in AML cells

Buoyed by these positive endogenous controls, we next examined the remaining CRISPRi hits to search for new biological insights and to generate hypotheses on the cellular mechanisms of decitabine action. First, we noted that the pathway-level analysis of our screen identifies mRNA processing pathways as a top-scoring enriched term (Supplementary Fig. 2f and Supplementary Table 2)^30,31^. Further analysis of these top hits revealed a strong enrichment for two specific RNA biological processes. Specifically, we observed that repression of RNA decapping enzymes such as *DCP1A*, *DCP2* and *DCPS* sensitizes HL-60 to decitabine (Fig.1c). We also observed that repression of multiple genes (*METTL3, YTHDF2, ZC3H13* and *CBLL1*) that regulate RNA methylation marks, specifically N^6^-methyladenosine or m^6^A, promoted resistance to decitabine. Together, these observations suggest that modulation of specific RNA regulatory pathways is a key determinant of response to DNA hypomethylation induced by decitabine. To independently validate the results from our screen, we chose 10 hit genes from our decitabine HL-60 CRISPRi screen (2 sgRNAs/gene) and used a mixed competition fluorescence cell survival CRISPRi knockdown assay to measure how perturbation of individual genes modulates response to decitabine. Our validation experiments demonstrated the reproducibility of our CRISPRi genome-scale screen measurements across all the resistance and sensitivity genes tested (Fig. 1d-f and Supplementary Fig. 2g). Interestingly, we observed that repression of *PTEN*, a tumor suppressor gene that is commonly mutated in cancer, sensitized HL-60 cells to decitabine (Fig. 1e).

We were intrigued by the connection between decitabine and RNA decapping quality control processes. To begin, we confirmed that repression of *DCP2* sensitizes cells to decitabine (Fig. 1e). We chemically validated that RNA decapping is a pro-survival dependency by combining RG3039, a clinical grade chemical inhibitor of *DCPS*, with decitabine^32,33^. We observed the combination of decitabine and RG3039 had synergistic anti-cancer activity *in vitro* in two independent AML cell models (Supplementary Fig. 3a-b). We also observed that the combination of decitabine and RG3039 synergistically induced caspase 3/7 activation and cell cycle arrest in HL-60 (Fig. 1g-h). Lastly, we profiled the transcriptional consequences of treating cells with DMSO, decitabine alone, RG3039 alone or decitabine and RG3039 together. Because previous literature has demonstrated HMAs can induce expression of endogenous retroviral elements, we mapped both protein coding transcript expression and ERV transcript expression. We observed that treatment with decitabine or RG3039 alone drives a transcriptional response, and that the combination of decitabine with RG3039 induces transcriptional responses shared with the single drug conditions but also drug combination specific transcriptional changes (Supplementary Fig. 3c-d). Gene ontology analysis comparing decitabine to decitabine plus RG3039 or DMSO to decitabine plus RG3039 demonstrated up regulation of term enrichment for biological processes such as myeloid differentiation and immune function, as well as down regulation for biological processes relating to methylation and protein translation (Supplementary Fig. 3e). For example, we observed the upregulation of positive regulators of TNFα cytokine production specifically in the decitabine plus RG3039 condition relative to decitabine alone (Fig. 1i). Additionally, we further examined myeloid differentiation as a top enriched process and observed broadly that treatment with decitabine or RG3039 alone induced a signature of differentiation relative to DMSO, and that this was further induced by the combined treatment of decitabine plus RG3039, suggesting that AML differentiation occurs from treatment with decitabine or RG3039 alone as well as in combination (Supplementary Fig. 3f-j). Lastly, prior studies have shown decitabine treatment alone can induce expression of atypical transcripts which in turn can induce an inflammatory response^10,34^. Our analysis of ERV transcriptional changes demonstrated that the combination of decitabine plus RG3039 strongly induced specific ERV transcripts, such as *LTR67B* (chr6:36350628−36351191), relative to DMSO or each single drug alone (Supplementary Fig. 3k-l). Notably, most ERVs do not change expression, and changes in expression are often not concordant across families or classes of ERVs. Together, this data suggests that RNA decapping is one of multiple processes which can affect response to decitabine in AML cells.

### Decitabine induces m^6^A hypermethylation of mRNA transcripts in AML cells

As highlighted above, we observed that repression of multiple genes encoding m^6^A methylation machinery promotes cellular resistance to decitabine (Fig. 1c,f). Top screen hits included the m^6^A-writer *METTL3*, the m^6^A-reader *YTHDF2* and the methyltransferase complex components *ZC3H13* and *CBLL1*. We validated that repression of *METTL3*, *YTHDF2*, and *ZC3H13* promotes resistance to DNMT inhibition by decitabine treatment in HL-60i over a time course using a mixed competition fluorescence cell survival CRISPRi knockdown assay (Fig. 2a). This result suggests regulation of RNA methylation modulates AML cell survival upon treatment with decitabine.

**Figure 2.**
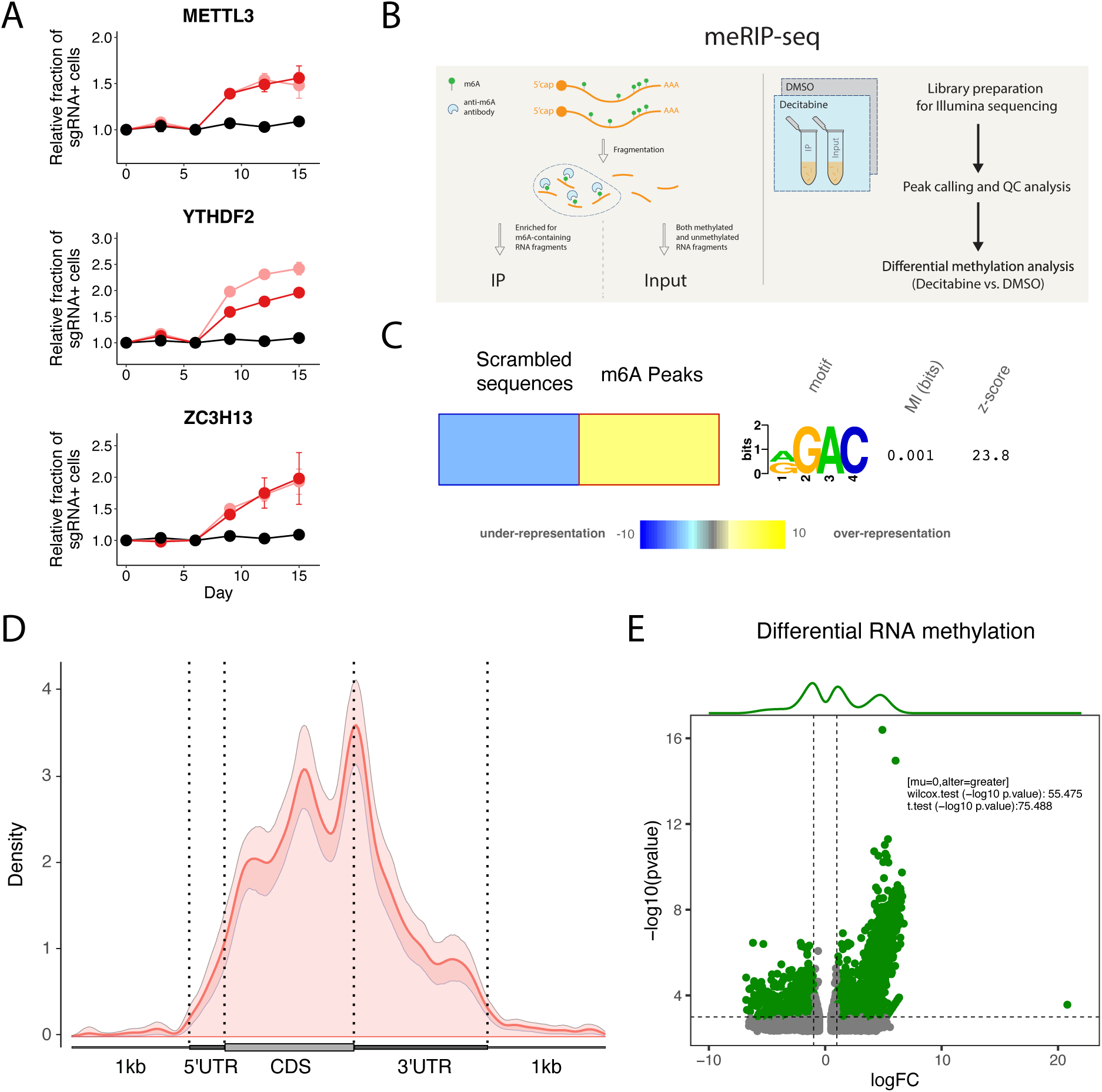
Decitabine treatment of HL-60 cells results in global m^6^A hypermethylation. (**a**) Validation of CRISPRi decitabine screen hits show that m^6^A-reader/writer complex genes promote resistance to decitabine treatment upon knockdown in HL-60i cells. HL-60i cells were transduced with a control sgRNA (black) or an active sgRNA (red or pink) and treated with DMSO or decitabine, and the proportion of sgRNA+ cells in the decitabine condition relative to DMSO was observed over time. Data are shown as means ± SD, two sgRNAs per gene and two replicates per sgRNA. (**b**) Schematic of MeRIP-seq experimental design and computational workflow. **(c)** The FIRE algorithm (in non-discovery mode) shows the known m^6^A motif RGAC ([AG]GAC) is enriched among predicted MeRIP-seq peaks relative to randomly generated sequences with similar dinucleotide frequencies. Data are shown as a heatmap, where yellow indicates over-representation and blue represents under-representation. Color intensity indicates the magnitude of enrichment. (**d**) Metagene plot shows distribution of m^6^A sites along transcripts with differential regional methylation and enrichment of m^6^A sites near the end codon. Transcripts are grouped into CDS (protein coding region), 5’ UTR (untranslated region) and 3’ UTR methylation based on the identified m^6^A sites. (**e**) Differential methylation analysis shows significant changes in RNA methylation peaks in HL-60 cells treated with decitabine (relative to DMSO). Peaks are called using the RADAR algorithm and visualized as annotated volcano plots. Wilcoxon and t-tests are used to assess statistical significance of global hypermethylation.

To systematically examine the molecular effect of decitabine treatment on m^6^A RNA methylation, we next performed methylated RNA immunoprecipitation sequencing (MeRIP-seq), a method for detection of m^6^A modifications (Fig. 2b)^35^. To assess the quality of this dataset, we first performed peak calling in control DMSO-treated samples followed by downstream analysis to recapitulate known features of the RNA modification sites across the transcriptome. We also performed a motif-enrichment analysis to ensure the enrichment of the RGAC ([AG]GAC) motif sequence, a known m^6^A motif, among predicted peaks (Fig. 2c)^36,37^. Finally, we confirmed the preferential localization of RNA methylation peaks near the stop codon, which is consistent with prior literature (Fig. 2d)^38^.

To then identify decitabine-induced hyper- and hypomethylated sites, we performed differential RNA methylation analysis to compare treatment with decitabine to DMSO controls^39^. Interestingly, we observed a significant increase in m^6^A RNA methylation peaks across mRNAs of protein coding genes upon decitabine treatment (Fig. 2e and Supplementary Table 3). Specifically, our analysis identified 2064 decitabine induced hypermethylated peaks (logFC >1 and p-value <0.005) but only 1399 hypomethylated peaks (logFC <−1 and p-value <0.005) (Supplementary Fig. 4b-d).

Additionally, it has been observed in AML cell lines and patient data that treatment with different HMAs such as decitabine induces transcriptional upregulation of different ERVs including retroposons, LINEs and SINEs^12,40,41^. It has also been shown that m^6^A RNA methylation regulates the levels of ERVs^42^. To evaluate the effect of decitabine treatment on ERV RNA methylation, we mapped our MeRIP-seq data to relevant annotations and followed similar analyses as discussed above to examine differential RNA methylation changes in ERVs^43^. Interestingly, we observed a significant enrichment of m^6^A methylation peaks across retroposon, LINE and SINE transcripts upon decitabine treatment (Supplementary Fig. 4e-f). Specifically, our analysis here identified 37, 180 and 131 hypermethylated peaks (logFC >1 and p-value <0.005) but only 9, 45 and 48 hypomethylated peaks (logFC <−1 and p-value <0.005) for retroposon, LINE and SINE transcripts, respectively.

Taken together, our findings suggest that treatment of AML cells with decitabine results in global CpG DNA hypomethylation along with a concomitant increase in m^6^A RNA methylation, and that HMA anti-cancer activity in AML cells may be modulated by genes that regulate m^6^A RNA methylation.

### A multiomics approach identifies genes regulated through m^6^A modifications

RNA methylation has been implicated in various aspects of the RNA life cycle in the cell, from RNA processing to RNA stability to translation, and more recently, crosstalk between epitranscriptome and epigenome^44–52^. To further understand the connection between global DNA hypomethylation and RNA dynamics in AML cells, we set out to interrogate, via an integrated multiomics approach, the effects of decitabine-induced RNA hypermethylation on AML cells. Here, we aimed to integrate comparisons between treatment with decitabine or DMSO from the following datasets: RNA-seq for differential gene expression and RNA stability, MeRIP-seq for RNA methylation, Ribo-seq for protein translation efficiency, and genome-scale CRISPRi functional genomics screening data. We first performed an RNA-seq time course experiment in the HL-60 AML model (Supplementary Fig. 5a) at 6, 72 and 120 hours following treatment with decitabine or DMSO. We used this data to perform differential gene expression analysis across conditions. We also used REMBRANDTS, a method we have previously developed for differential RNA stability analysis, to estimate post-transcriptional modulations in relative RNA decay rates (Fig. 3a-b)^53–58^. We performed gene set enrichment analysis of differential mRNA stability and expression across all three time points for the HL-60 cell line (Supplementary Fig. 5b-c)^59^. For expression, we observed enrichment for largely expected ontologies, such as immune receptor activity and regulation of cell killing^10,12,14,34^. Interestingly, for post-transcriptional modulations in RNA stability, we observed previously unexplored terms, such as sterol biology. Moreover, to also capture patient heterogeneity, we performed RNA-seq on a panel of five additional AML cell lines treated with decitabine or DMSO. Across all six AML cell lines, we observed that decitabine treatment induced widespread changes in RNA transcript abundance and RNA stability with varying degrees of concordant RNA expression and stability changes (Fig. 3c-d).

**Figure 3.**
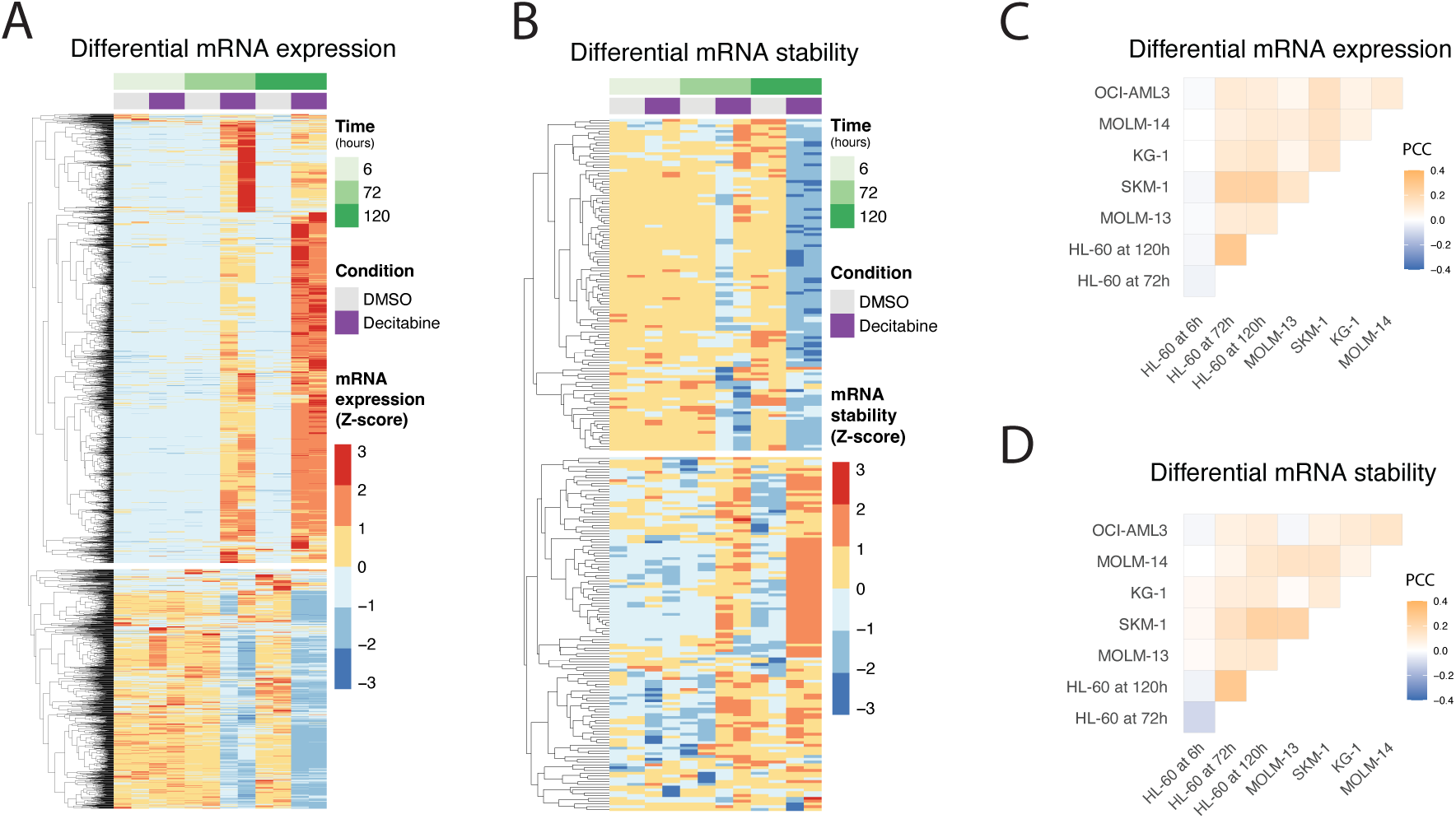
An analysis of differential gene expression and RNA stability across multiple AML cell lines and time points following decitabine treatment. (**a-b**) RNA-seq reveals genes with significant changes in **(a)** gene expression and **(b)** RNA stability in HL-60 cells following treatment with decitabine vs. DMSO. Data are shown as heatmaps displaying counts (of two replicates) row-normalized into Z-scores, grouped by treatment condition and time. Differential RNA expression was calculated using our Salmon-tximport-DESeq2 pipeline. RNA stability was predicted using the REMBRANDTS algorithm and differential RNA stability was calculated using limma. (**c-d**) RNA-seq shows varying degrees of concordance of differential **(c)** gene expression and **(d)** RNA stability across a panel of six AML cell lines. The correlation analysis was performed on the logFC values from (**c**) DESeq2 and (**d**) limma results for cells treated with decitabine vs. DMSO. Data are shown as correlation matrices with Pearson’s correlation coefficients (PCC).

Given that RNA m^6^A methylation marks have been previously implicated in translational control, we used Ribo-seq to measure changes in the translational efficiency landscape of HL-60 cells treated with decitabine or DMSO^47,60^. Treatment with decitabine had little effect on translation efficiency, and we did not observe a concerted change in the translation efficiency of hypermethylated mRNAs (Supplementary Fig. 6a-d). In other words, changes in translation efficiency of mRNAs that are differentially methylated in decitabine-treated cells are not likely to be responsible for cellular sensitivity to this drug.

Having ruled out translational control as the mechanism through which RNA methylation may be involved, we next sought to identify genes whose RNA hypermethylation drives cellular sensitivity to decitabine through other post-transcriptional regulatory programs. Since m^6^A RNA methylation has been shown to reduce RNA stability and expression, we intersected our set of decitabine-induced hypermethylated genes with those that are downregulated in decitabine treated cells, and their lower expression is associated with higher sensitivity to decitabine in our functional CRISPRi screen^61^. In this analysis, we observed ten genes that were sensitizing hits in the CRISPRi screen and upon decitabine treatment, showed RNA hypermethylated peaks and lower mRNA levels (Fig. 4a-b). We observed that these genes collectively regulate nuclear processes (*INTS5*, *INO80D*, *ZNF777*, *MYBBP1A*, *RNF126*, *RBM14-RBM4*)) or metabolism (*SQLE*, *DHODH*, *PMPCA*, *SLC7A6*). From this list we selected *SQLE* and *INTS5* and first validated that repression of each gene by CRISPRi conferred sensitivity to decitabine treatment in HL-60 cells (Fig. 4c). We then validated that their mRNA abundance is decreased and m^6^A methylation is increased following decitabine treatment (Fig. 4d and Supplementary Fig. 7a-b). Consistently, we observed that *SQLE* and *INTS5* pre-mRNA levels do not change, showing that the decreased mRNA levels are not due to a decrease in transcription. Additionally, we further examined mRNA stability of each gene in decitabine-treated cells by using α-amanitin to inhibit RNA polymerase II and observed that mRNA decay rates were significantly higher upon decitabine treatment (Fig. 4e). Lastly, we were intrigued by whether the increase in m^6^A methylation from decitabine occurred through *METTL3* given the methyltransferase’s direct role in regulating m^6^A methylation. Interestingly, we observed that upon *METTL3* knockdown, decitabine treatment no longer resulted in a significant increase in m^6^A methylation, suggesting that the decitabine-induced hypermethylation of these transcripts occurs through *METTL3* (Fig. 4f). These results together suggest that we have identified a small number of mRNAs that are downregulated upon decitabine treatment, likely through post-transcriptional processes including increased m^6^A methylation that is mediated by *METTL3*, and that these genes may be functionally important for cellular response to decitabine.

**Figure 4.**
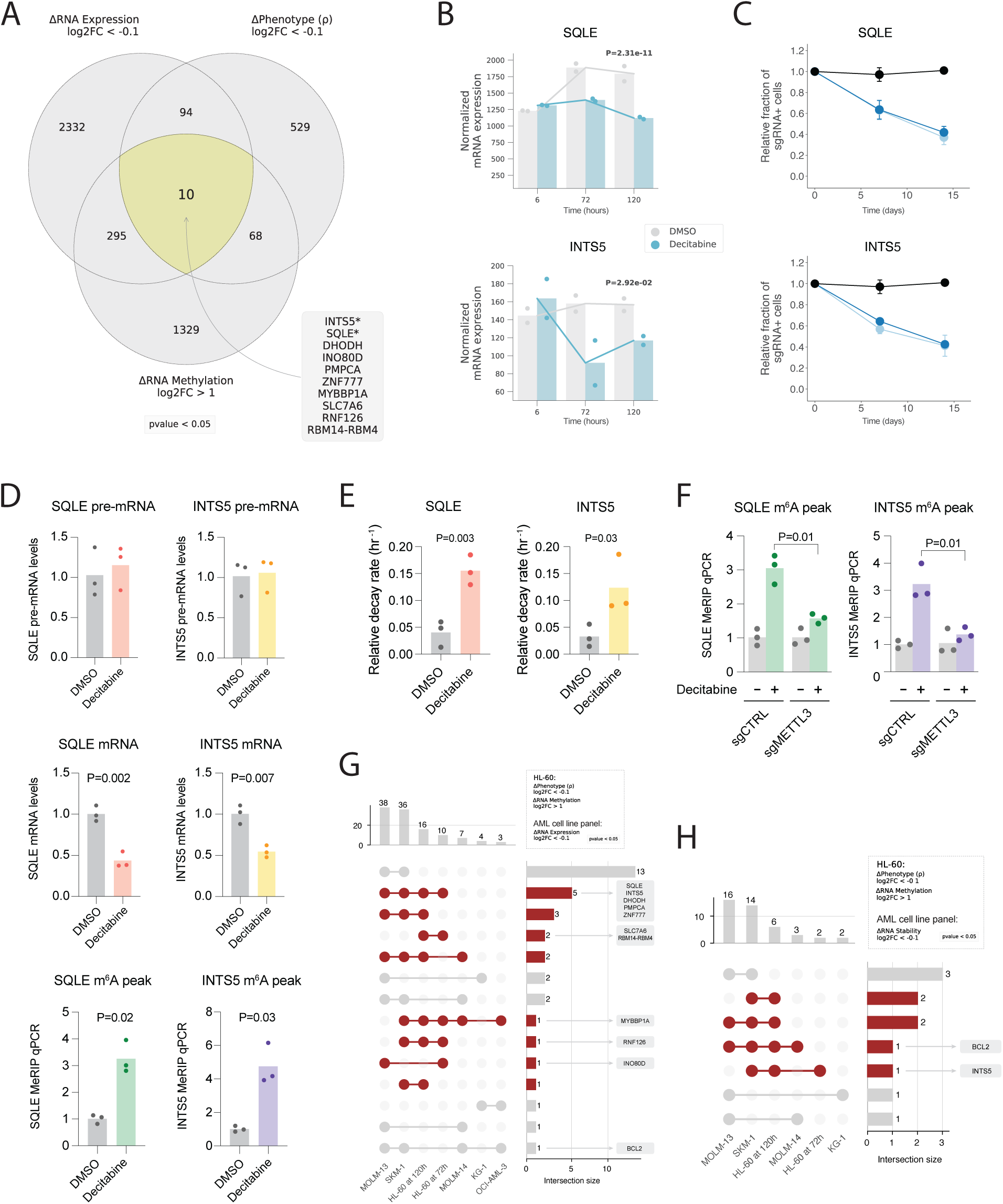
Charting genes likely downregulated due to m^6^A hypermethylation in HL-60 cells treated with decitabine and validating *SQLE* and *INTS5*. (**a**) Venn diagram visualization of three sets of genes across multiomics datasets (i.e., CRISPRi screen, RNA-seq and MeRIP-seq) for HL-60 cells treated with decitabine vs. DMSO. 10 overlapping genes were shown to have (1) a sensitizing phenotype in our CRISPRi screen, (2) RNA hypermethylation upon decitabine treatment and (3) downregulation of mRNA upon decitabine treatment. **(b)** Normalized RNA-seq counts for *SQLE* and *INTS5* in HL-60 cells treated with decitabine vs. DMSO at 6 hours, 72 hours and 120 hours. Data are shown as two replicates and p-values were generated using a likelihood ratio test in DESeq2 comparing the decitabine and DMSO conditions at 72 hours. (**c**) Validation of CRISPRi decitabine screen hits show that *SQLE* and *INTS5* knockdown promotes sensitivity to decitabine treatment in HL-60i cells. HL-60i cells were transduced with a control sgRNA (black) or an active sgRNA (blue) and treated with DMSO or decitabine, and the proportion of sgRNA+ cells in the decitabine condition relative to DMSO was observed over time. Data are shown as means ± SD, two sgRNAs per gene and two replicates per sgRNA. (**d**) MeRIP-RT-qPCR in HL-60 cells treated with DMSO (gray) or decitabine (colored) validates decitabine-induced mRNA decay and RNA hypermethylation of *SQLE* and *INTS5* transcripts. Three sets of primers were designed to capture abundances of pre-mRNA (top), mature mRNA (middle) and predicted m^6^A hypermethylated loci for each gene (bottom). Data are shown as three replicates and one-tailed Mann-Whitney U-tests were used to assess statistical significance. (**e**) RT-qPCR validation of decitabine-induced mRNA decay of *SQLE* and *INTS5* using α-amanitin. HL-60 cells were treated with DMSO (gray) or decitabine (colored) ± α-amanitin and RT-qPCR captured mRNA abundance. Relative decay was defined as the ratio between samples with and without α-amanitin for each respective condition. Data are shown as three replicates, and one-tailed Mann-Whitney U-tests were used to assess statistical significance. (**f**) MeRIP-RT-qPCR in HL-60 cells reveals *METTL3* as a regulator of decitabine-induced m^6^A hypermethylation of *SQLE* and *INTS5*. Cells were transduced with a control sgRNA or *METTL3*-targeting sgRNA, treated with DMSO (gray) or decitabine (colored), and MeRIP-RT-qPCR captured abundance of predicted m^6^A hypermethylated loci. Data are shown as three replicates and one-tailed Mann-Whitney U-tests were used to assess statistical significance. (**g-h**) UpSet plots visualizing the intersection between genes which were (1) RNA hypermethylated upon decitabine treatment in HL-60 and (2) sensitizing hits in the HL-60 CRISPRi screen with (**g**) genes downregulated and (**h**) RNA destabilized across six AML cell lines.

To extend our observations, we also identified genes that (i) were downregulated upon decitabine treatment across our panel of six AML cell lines, (ii) sensitizing hits in our HL-60 CRISPRi screen, and (iii) showed hypermethylated peaks upon decitabine treatment in our MeRIP-seq HL-60 data (Fig. 4g-h). Although this analysis converges on a very small number of genes, we were nevertheless intrigued by the possibility that several nominated genes could serve as a link between RNA methylation and the cell death induced by decitabine.

### Comparative CRISPRi functional genomics experiments reveal common and specific genes modulating cellular response to decitabine in additional AML models

Given the known heterogeneity of AML, we chose to perform genome-scale CRISPRi screens in two additional AML models to further examine the degree of common and specific mechanisms across cell lines that regulate cellular response to decitabine. For this we used SKM-1 and MOLM-13 cells, which are established models of AML. Comparing the known driver mutations in these AML models, we noted that SKM-1 is *TP53* and *KRAS* mutant, which similarly to HL-60, captures the biology of high-risk AML and more generally of an aggressive human cancer. Meanwhile, MOLM-13 is *FLT3*-ITD and *MLL*-fusion but *TP53* wild-type. We also examined the genetic status of the RNA-related genes of interest from our HL-60 screen and noted that these genes are not commonly mutated across AML (Supplementary Fig. 8a-b). We engineered CRISPRi cell lines for each model and performed genome-scale CRISPRi screens to identify genes that regulate response to decitabine (∼IC30; 15-100 nM) as described above and compared the results with the HL-60 screen (Supplementary Fig. 8c-f).

Similar to the HL-60 screen, we observed that the SKM-1 and MOLM-13 screens also captured mRNA processing as an enriched term across top hits and positive control genes whose knockdown is known to impact drug resistance, namely *DCK*, *SLC29A1* and *DCTD* (Supplementary Fig. 8d-f and Supplementary Tables 4-5)^18,19,29^. Additionally, we observed that repression of *METTL3* promoted resistance to decitabine across all three cell lines. As expected from the heterogeneity of AML, we also observed differences across cell lines with respect to genes that modulate response to decitabine. Interestingly, the two cell lines classified as *TP53*-inactive (HL-60 and SKM-1), and are representative of the high-risk AML patient cohort that benefits from the combination therapy of decitabine and venetoclax, revealed *BCL2* and *MCL1* as sensitizing hits in the presence of decitabine, while the *TP53*-wild-type cell line (MOLM-13) did not^20-21^. Additionally, repression of genes encoding RNA decapping enzymes such as *DCP2* and *DCPS* sensitized HL-60 and SKM-1 cells, but not MOLM-13 cells, to decitabine treatment.

In summary, comparison of genome-scale decitabine CRISPRi screens in three AML models reveals common and unique regulators of response. These findings are in line with our understanding of the heterogeneity of AML biology and suggest that therapeutic strategies in AML should be evaluated in multiple models representative of diverse tumors.

## DISCUSSION

Our experiments identify previously known and unknown genes and pathways that modulate cellular response to decitabine, a clinically approved HMA with poorly understood cellular mechanisms of action. Our results unexpectedly reveal a key role for RNA dynamics in modulating the response to DNA hypomethylation induced by decitabine.

Specifically, we observed that genes which are thought to regulate mRNA decapping promote cellular resistance to decitabine. One hypothesis for why loss of RNA decapping enzyme activity sensitizes AML cells to decitabine is that this RNA quality control pathway becomes an induced dependency upon decitabine treatment due to repressed or aberrant transcripts that accumulate upon decitabine-induced DNA hypomethylation. Alternatively, some RNA decapping proteins are also key regulators of splicing, so it may be that this biology is more complex with respect to transcription than currently appreciated^62,63^.

We also found that genes responsible for writing and reading m^6^A RNA methylation mediate cellular response to decitabine. While emerging evidence suggests potential cellular crosstalk between DNA and RNA methylation, the direct connection between the two processes, particularly in the context of m^6^A RNA methylation and DNMT inhibitors, remains underexplored^50,52,64^. Our results demonstrate that decitabine treatment induces global m^6^A hypermethylation in AML cells, and that inhibition of a key adenosine methyltransferase *METTL3* promotes resistance to decitabine. Given that *METTL3* has been previously shown to be a potential therapeutic vulnerability in AML^65,66^, it is intriguing to posit why its inhibition may promote resistance to a drug used in clinic to treat high-risk AML. Given all known human methyltransferase enzymes use S-adenosyl methionine (SAM) as a cofactor for transfer of methyl groups, one hypothesis arises in which treatment of cells with decitabine results in global inhibition of DNMTs, resulting in increased SAM levels and subsequently hypermethylation of mRNAs leading to transcript instability and cell death. To our knowledge, crosstalk between methyltransferase enzymes and different macromolecular substrates is not known, and this hypothesis may merit further investigation.

Our efforts may have several translational implications for AML patients who are treated with decitabine. First, we experimentally confirm that decitabine induces *TP53* independent apoptosis in experimental models. In line with this, our results genetically re-nominate a clinically efficacious combination therapy of decitabine and a *BCL2* inhibitor, which together likely induces synergistic apoptosis^20-21^. We also demonstrate through both genetic and chemical approaches that RNA decapping pathways promote the survival of AML cells treated with decitabine *in vitro*. Lastly, we observe dysregulation of specific transcripts that may have therapeutic relevance, such as *SQLE*, where studies in various cancer models have suggested that its inhibition may suppress tumor growth, or *DHODH*, which has previously been implicated in AML and currently has an inhibitor in clinical trials for relapsed/refractory AML^67–71^.

We anticipate that our study serves as an integrated multiomics resource for understanding AML cellular response to decitabine and nominates new connections between cell death, DNA methylation and RNA dynamics.

## Data Availability

The data that support the findings of this study are openly available in NCBI Gene Expression Omnibus (GEO) with reference number GSE222886 (RNA-seq, meRIP-seq, Ribo-seq).

## Acknowledgments

We would like to thank members of the Goodarzi and Gilbert labs for helpful discussions and assistance, including Dr. Gary Wang, Dr. Becky Xu Hua Fu, Dr. Benedict Choi, Laine Goudy, and Tanvi Joshi. We also thank Dr. Felix Krueger for helping A.A. troubleshoot Bismark. L.A.G. is funded by an NIH New Innovator Award (DP2 CA239597), a Pew-Stewart Scholars for Cancer Research award and the Goldberg-Benioff Endowed Professorship in Prostate Cancer Translational Biology. H.G. is an Era of Hope Scholar (W81XWH-2210121) and supported by R01CA240984 and R01CA244634. This work is also supported by the Laboratory of Genomics Research established by GSK, UCSF and UC Berkeley. Sequencing was performed at the UCSF CAT, supported by UCSF PBBR, RRP IMIA and NIH 1S10OD028511-01 grants.

## Author Contributions

A.Y.G., A.A., R.D., L.P., H.G. and L.A.G. conceived and designed the experiments. A.Y.G., A.A., R.D., A.N., L.F., K.G., J.G., S.L., K.K., M.B.P., L.P. and H.G. performed the experiments. A.Y.G., A.A., R.D., H.A., M.T.M., L.P., H.G. and L.A.G. analyzed the screen and screen validation results. A.A. and H.G. integrated and analyzed the multiomics datasets. A.Y.G., A.A., R.D., H.G. and L.A.G. prepared the manuscript.

## METHODS

### Cell culture and reagents

HL-60 and KG-1 cells were obtained from the American Type Culture Collection. SKM-1, MOLM-13 and OCI-AML3 cells were obtained from the Leibniz Institute DSMZ (German Collection of Microorganisms and Cell Cultures). MOLM-14 cells were obtained from the Shannon Lab at the University of California, San Francisco (UCSF). HEK-293T cells were obtained from the Weissman Lab at UCSF. HL-60, OCI-AML3 and KG-1 cells were cultured in Iscove’s Modified Dulbecco’s Medium (Gibco) supplemented with 20% fetal bovine serum (Seradigm), 100 U/mL penicillin (Gibco), 100 ug/mL streptomycin (Gibco) and 0.292 mg/mL glutamine (Gibco). SKM-1, MOLM-13 and MOLM-14 cells were cultured in RPMI-1640 medium (Gibco) supplemented with 20% FBS, penicillin, streptomycin and glutamine. HEK-293T cells were cultured in Dulbecco’s Modified Eagle Medium (Gibco) supplemented with 10% FBS and penicillin, streptomycin and glutamine. All cell lines were grown at 37℃ and 5% CO_2_ and were tested for mycoplasma contamination using the MycoAlert PLUS Mycoplasma Testing Kit (Lonza) according to the manufacturer’s instructions.

Decitabine powder was obtained from Selleck Chemicals and stored at -20℃. A stock solution of decitabine was created by reconstituting decitabine powder in dimethyl sulfoxide (DMSO) at a final concentration of 10 mM. The stock solution was aliquoted and stored at -80℃ until experimental use. RG3039 and α-amanitin were obtained from MedChemExpress.

### DNA transfections and lentivirus production

HEK-293T cells were transfected with pMD2.G, pCMV-dR8.91 and a transfer plasmid using the TransIT-LT1 Transfection Reagent (Mirus Bio) and 8 ng/uL polybrene. Culture medium was exchanged with fresh medium supplemented with ViralBoost (Alstem) one day post-transfection. Lentiviral supernatant was collected, filtered through a 0.44 μm filter (Millipore) and used fresh (for CRISPRi screening) or concentrated via ultracentrifugation at 25,000 rpm for 90 minutes and frozen (for all other methods) three days post-transfection.

### CRISPRi screen

#### CRISPRi cell line generation

HL-60 cells were transduced with Ef1a-dCas9-BFP-KRAB and sorted twice for BFP positive cells on a BD FACS Aria III. Sorted cells were diluted to single cell concentration (5, 1 or 0.2 cells per well) and plated into 96-well plates. Individual clones were expanded and assayed for CRISPRi activity by transducing sgRNAs targeting five essential genes (PLK1, HSPA9, AARS, POLR1D, DNAJC19) and assessing for relative depletion of GFP (i.e., sgRNA positive cells) via flow cytometry between day 3 and day 9 post-transfection. The clone with the highest relative GFP depletion was selected to be the HL-60 CRISPRi cell line for downstream experiments. SKM-1 and MOLM-13 cells were transduced with Ef1a-dCas9-BFP-KRAB and sorted twice for BFP positive cells on a BD FACS Aria III. Cells were then assayed for CRISPRi activity by transducing sgRNAs targeting two essential genes (PLK1, HSPA9) and assessing for relative depletion of GFP (i.e., sgRNA positive cells) via flow cytometry between day 3 and day 9 post-transfection.

#### CRISPRi screen experimental procedure

Genome-scale CRISPRi screens were performed similarly to those previously described^23^. The human CRISPRi-v2 sgRNA library (top 5 sgRNAs per gene) was transduced into HL-60, SKM-1 and MOLM-13 cells at 250 to 500-fold coverage. Cells were resuspended in lentiviral supernatant with 8 μg/mL polybrene in 6-well plates and centrifuged at 1000 *g* for 2 hours at room temperature. Cells were resuspended into fresh medium following spinfection. 72 hours following infection, cells were seeded at 1,000,000 cells/mL for puromycin selection (0.5-1 ug/mL). Following puromycin selection, “time-zero” samples were harvested at 500x library coverage. The remaining cells were divided into two conditions, DMSO and decitabine, two replicates per condition. For the decitabine condition, cells were treated with decitabine at low dose (∼IC30; 15-100 nM) every 24 hours for 72 hours. For HL-60, cells were cultured in static T150 flasks (Corning) and split when appropriate while maintaining 500x coverage; after 19 days of growth, cells were harvested at 500x coverage. For SKM-1 and MOLM-13, cells were cultured in 250 mL OptimumGrowth (Thomson) shaking flasks with a shaking speed of 120 rpm and split when appropriate while maintaining a minimum coverage of 500x; after 12 days of growth, cells were harvested at 500-1000x coverage. Genomic DNA was isolated from all samples and the sgRNA-encoding region was enriched, amplified and processed for sequencing on the Illumina HiSeq 4000 (50 base pair single end reads) as previously described^72^.

#### CRISPRi screen computational analysis

Sequencing reads were trimmed, aligned to the human CRISPRi-v2 sgRNA library and counted using a previously described pipeline (https://github.com/mhorlbeck/ScreenProcessing). Growth (γ) and drug sensitivity/resistance (ρ) phenotypes were calculated based on sgRNA frequencies across conditions^23^. Gene phenotypes were calculated by taking the mean of the top three sgRNA phenotypes per gene by magnitude. Gene phenotype p-values were calculated using the Mann-Whitney test comparing the gene-targeting sgRNAs with a set of non-targeting control sgRNAs. For genes with multiple annotated transcription start sites (TSS), sgRNAs were first clustered by TSS, and the TSS with the smallest Mann-Whitney p-value was used to represent the gene. Hits were defined as genes with a phenotype Z-score greater or equal to 6. Z-scores were calculated by dividing the gene phenotype by the standard deviation of the non-targeting sgRNA phenotypes^23^.

To assess pathway-level enrichment of gene phenotypes in the CRISPRi screen, we used blitzGSEA, a Python package for the computation of Gene Set Enrichment Analysis (GSEA) (https://github.com/MaayanLab/blitzgsea)^30^. We obtained gene ontology (GO) gene sets from MSigDB (version 7.4.) and then conducted two separate analyses: (1) To identify smaller, focused pathways associated with drug sensitivity or resistance, we performed GSEA analysis on genes ranked by ρ phenotype and defined minimum and maximum thresholds for gene set size when running the ‘gseà function (‘min_size=15’ and ‘max_size=150’)^31,73^. Thus, positive normalized enrichment scores (NES) corresponded to gene sets enriched among positive ρ phenotypes (i.e., resistance phenotypes) and negative NES corresponded to gene sets enriched among negative ρ phenotypes (i.e., sensitivity phenotypes). (2) To identify broader pathways associated with drug response irrespective of ρ phenotype direction, we performed GSEA analysis on genes ranked by 1 – Mann-Whitney p-value (calculated for each ρ phenotype as above) and set a minimum threshold for gene set size (i.e., ‘min_size=200’).

### Individual sgRNA validation

Individual sgRNAs were validated using a competitive growth assay as previously described^23^. Briefly, sgRNA protospacers with flanking BstXI and BlpI restriction sites were cloned into the BstXI/BlpI-digested pCRISPRia-v2 plasmid (Addgene #84832). Protospacer sequences are listed in Supplementary Table 1. Individual sgRNA vectors (including a non-targeting control sgRNA) were then packaged into lentivirus as described above and transduced into HL-60 CRISPRi cells in duplicate. Three days after transduction, cells were treated with DMSO or 100 nM decitabine. The proportion of sgRNA-expressing cells was measured by flow cytometry on an LSR II (BD Biosciences) gating for GFP expression. The individual sgRNA phenotype was calculated by dividing the fraction of sgRNA-expressing cells in the treated condition by the fraction of sgRNA-expressing cells in the untreated condition. To confirm gene knockdown at the transcriptional level, mRNA abundances were measured in CRISPRi cells transduced with gene-targeting sgRNAs and were quantified relative to mRNA abundances in cells transduced with a non-targeting control sgRNA, as previously described^74^.

### Reanalysis of public bisulfite sequencing data in HL-60 cells

Shareef et. al, as part of a study to introduce their extended-representation bisulfite sequencing method, treated HL-60 cells with DMSO (GSM4518676) or 300 nM decitabine (GSM4518677) and harvested cells after 5 days^25^. Raw FASTQ files were downloaded using the SRA Toolkit. TrimGalore and Bismark were used to preprocess and map bisulfite-treated reads to the h38 reference genome and subsequently call cytosine methylation^75^. We followed the Bismark standard pipeline, which includes four functions: (1) ‘bismark_genome_preparation’, (2) ‘bismark’, (3) ‘deduplicate_bismark’ and (4) ‘bismark_methylation_extractor’ which extracts context dependent (CpG/CHG/CHH) methylation.

Differential CpG DNA methylation analysis was performed using the methylKit R package^76^. CpG methylation data from Bismark was imported and the ‘getMethylationStats’ function was used to calculate descriptive statistics. To search for differentially methylated tiles, the ‘tileMethylCounts’ function was used with options ‘win.size=1000’ and ‘step.size=1000’. Differentially methylated regions (DMRs) scored by % methylation difference and q-value were calculated using the ‘calculateDiffMeth’ function. A one-sample, one-sided (lower-tail) t-test was used to test for statistically significant global DNA hypomethylation.

### DCPS and RG3039 drug synergy experiments

#### Cell viability assay and Bliss excess score calculation

Cells were seeded into 96-well plates at 100,000 cells/mL in duplicate and were treated with decitabine (seven-point 1:3 dilution series from 0.5 uM to 0.002 uM), RG3039 (seven-point 1:4 dilution series from 10 uM to 0.010 uM) or the combination of both drugs at all possible dose combinations. Control cells treated with DMSO were counted at day 3, and all cells were split at the ratio required to dilute control cells to a concentration of 100,000 cells/mL. Raw fluorescence units (RLUs) were assessed at day 3, day 5 and day 7 for each condition using the CellTiter-Glo (CTG) luminescence-based assay (Promega). Diluted CTG reagent (100 uL 1:4 CTG reagent to PBS) was added to cells (100 uL) and the mixture was pipetted up and down to ensure complete cell lysis. Luminescence was then assayed using a GloMax Veritas Luminometer (Promega). To calculate the proportion of viable cells, RLUs from the CTG assay were averaged between replicates and normalized to the DMSO control. The proportion of inhibited cells was calculated as one minus the proportion of viable cells. Drug synergy was determined by calculating the Bliss excess score (Bliss 1956)^77^, i.e.

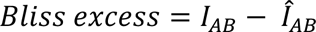

where *I*_*AB*_ represents the observed proportion of inhibited cells at drug doses *A* and *B* and 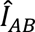 represents the expected proportion of inhibited cells assuming Bliss independence, i.e.

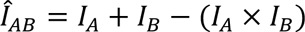

#### Cleaved caspase 3/7 assay

Cells were seeded into 24-well plates at 100,000 cells/mL in triplicate and were treated with decitabine (50 nM, 100 nM or 200 nM on days 0, 1 and 2) with and without RG3039 (2 uM on day 0). Cells were harvested on day 5 and the proportion of apoptotic cells was assessed using the NucView 488 Caspase-3 Assay Kit (Biotium) according to the manufacturer’s instructions and an Attune NxT flow cytometer (Thermo Fisher Scientific) gating on the BL-1 channel.

#### Cell cycle assay

Cells were seeded into 24-well plates at 100,000 cells/mL in triplicate and were treated with decitabine (50 nM, 100 nM or 200 nM on days 0, 1 and 2) with and without RG3039 (2 uM on day 0). Cells (500,000–1,000,000 per sample) were harvested on day 5 and the proportions of cells in each phase of the cell cycle were assessed using the FxCycle Violet Kit (Thermo Fisher Scientific) and an Attune NxT flow cytometer (Thermo Fisher Scientific) gating on the VL-1 channel. Briefly, cells were washed once with PBS, fixed with 70% ethanol overnight at -20 °C, pelleted, and then washed with PBS 1–2 times. Cells were then resuspended in 1 mL permeabilization buffer (PBS with 1% FBS and 0.1% Triton X-100) and 1 uL Fx cycle and stained for 30 minutes in the dark before being analyzed via flow cytometry.

### RNA-seq experimental procedures

#### 3’ RNA-seq

3’ RNA-seq was performed to assess differential gene expression following decitabine and/or RG3039 treatment. Cells were seeded into 6-well plates at 100,000 cells/mL in duplicate and were treated with decitabine (100 nM on days 0, 1 and 2), RG3039 (2 uM on day 0), both drugs or DMSO. On day 3, RNA was extracted using the RNeasy Mini Kit (Qiagen) according to the manufacturer’s instructions. RNA-seq libraries were prepared using the QuantSeq 3′ mRNA-Seq Library Prep Kit FWD for Illumina (Lexogen) and assessed on a BioAnalyzer 2100 (Agilent) for library quantification and quality control. RNA-seq libraries were sequenced on an Illumina HiSeq 4000 using single-end, 50–base pair sequencing.

#### Stranded RNA-seq

Stranded RNA-seq was performed for experiments in which strand directionality was required for downstream analysis. Cells were seeded into 6-well plates at 100,000 cells/mL in duplicate or triplicate and were treated with decitabine (100 nM on days 0, 1 and 2) or DMSO. At 6, 72 and/or 120 hours, RNA was extracted using the RNeasy Mini Kit (Qiagen) according to the manufacturer’s instructions. For HL-60 experiments, RNA-seq libraries were prepared using the ScriptSeq v2 kit (EpiCentre). Total RNA was depleted using RiboZero Gold (EpiCentre) and purified using the MinElute RNA kit (Qiagen). For all other cell lines, RNA-seq libraries were prepared using the SMARTer Stranded Total RNA Sample Prep Kit - HI Mammalian kit (Takara) due to retirement of the ScriptSeq v2 kit. Total RNA was depleted using the RiboGone module included with the SMARTer kit. All RNA-seq libraries were assessed on a BioAnalyzer 2100 (Agilent) for library quantification and quality control and sequenced on an Illumina HiSeq 4000 using single-end, 50– base pair sequencing.

### Differential gene expression analysis

#### The Salmon-tximport-DESeq2 pipeline

We used a workflow hereafter referred to as the “Salmon-tximport-DESeq2 pipeline” to perform differential gene expression analysis. Salmon (version 1.2.1) was first used to quantify transcript abundance^55^. A Salmon index was generated using the GENCODE (version 34) genome annotation, and subsequently the ‘salmon quant’ tool was used with the ‘--validateMappings’ option to calculate transcript abundances^78^. Then, the R package tximport was used to import Salmon results into R and perform data preparation^54^. The ‘summarizeToGenè function was used to collapse transcript abundances to the gene level. From here, the R package DESeq2 was used for differential gene expression analysis^53^. We first extracted normalized counts for each RNA-seq experiment using DESeq2 by running the ‘estimateSizeFactors’ function and then the ‘counts’ function with option ‘normalized=TRUÈ. For each individual experiment, the DESeq2 statistical model was modified based on the experimental design. For experimental designs with multiple variables (e.g., multiple drug conditions, time points, etc.), we used the likelihood ratio test (LRT) to perform differential expression analysis. The LRT is conceptually similar to an analysis of variance (ANOVA) calculation in a linear regression model^79^. In these cases, we specified the model design in the ‘DESeq2’ function as ‘∼0 + variable1 + variable2 + variable1:variable2’ and the option ‘test=LRT’. In simple experimental designs with one variable (e.g., DMSO vs. decitabine treatment), DESeq2 was used with default options (i.e., a Wald test was used instead of a LRT). In these cases, the model design was specified as ‘∼cond’. For experiments with batch effects, the model design was specified as ‘∼cond + reps’.

### Differential RNA stability analysis

#### The STAR-featureCounts-REMBRANDTS-limma pipeline

For analyses which required measurements of pre-mRNA and mature mRNA abundances from RNA-seq samples (i.e., differential RNA stability analysis), we used a workflow hereafter referred to as the “STAR-featureCounts-REMBRANDTS-limma pipeline”. RNA-seq sequencing reads were first aligned to the hg38 reference genome using STAR (version 2.7.3a)^56^. Then, featureCounts was used to quantify intron and exon level counts. Finally, REMBRANDTS was used to calculate mRNA stability as previously described (https://github.com/csglab/REMBRANDTS)^58^. Briefly, the package estimates a gene-specific bias function that is subtracted from Δexon–Δintron calculations to provide unbiased mRNA stability measurements. To assess differential RNA stability changes, we used limma, which was designed for microarray experiments and serves a similar function to DESeq2, though it supports negative values (relevant for RNA stability analysis)^57^. The model designs used here are analogous to the designs for differential expression analysis described above.

### Gene set enrichment analysis using PAGE algorithm

Briefly, PAGE quantizes differential measurements into equally populated bins and then, for every given geneset, calculates the mutual information (MI) between each cluster bin and a binary vector of pathway memberships for genes in a given gene set^59^. The significance of each MI value is then assessed through a randomization-based statistical test and hypergeometric distribution to determine whether there is over or under representation of a gene set in each cluster bin. The final result is a p-values matrix in which rows are gene sets and columns are cluster bins (visualized as heatmaps). Code for iPAGE and onePAGE analyses are available at https://github.com/abearab/pager.

#### iPAGE run for MSigDB gene sets

The iPAGE algorithm was used for gene set and pathway enrichment analysis on differential RNA expression and stability results^59^. MSigDB (version 7.4.) was downloaded and modified to be compatible with iPAGE workflow^73^. iPAGE was used in continuous mode, which accepts gene-level numeric values (e.g., logFCs) as input.

#### onePAGE run for single gene set analysis

For a selected list of genes, the PAGE run is performed on the single gene set as first input and gene-level numeric values (e.g., log fold changes) as the other input – this form of the analysis is called onePAGE. This analysis applied to a specific gene set for multiple inputs (e.g., differentially expressed genes from different conditions) and results shown as heatmap where each row corresponds to an input condition and each column corresponds to a cluster bin.

### Pre-processing HERV annotations for alignment tasks

Annotations in BED12 format were downloaded from the Human Endogenous RetroViruses Database^43^. To prepare these annotations for alignment tasks, i.e., building Salmon and STAR indices, CGAT Apps was used to convert BED12 files to GTF format (‘cgat bed2gff --as-gtf’) and the ‘getfastà module from bedtools (with options ‘-name+ -split’) was used to convert BED12 files to FASTA format^80,81^. Reproducible scripts for preparing ERV annotations for alignment tasks are available at https://github.com/abearab/HERVs.

### RNA-seq workflows for specific experiments

#### Decitabine and RG3039 drug combination experiments

We performed 3’ RNA-seq on HL-60 cells treated with DMSO, decitabine alone, RG3039 alone or both drugs for 72 hours in duplicate (see above for experimental procedures). Raw sequencing data were processed using our Salmon-tximport-DESeq2 pipeline (see above). DESeq2 was used to conduct differential gene expression analysis using a likelihood ratio test and the model design ‘∼0 + decitabine + rg3039 + decitabine:rg3039’. Pathway enrichment was assessed using iPAGE (see above). For PCA analysis, the ‘varianceStabilizingTransformation’ function from the DESeq2 package was used to prepare counts. The ‘plotPCÀ function was used to calculate PC variances as percentages. Finally, ‘ggplot2’ was used to visualize a two-dimensional representation of the PCA analysis. Bar plots were used to visualize mRNA abundances (measured as log2 of transcripts per million [TPM]) of differentiation markers across conditions. Gene set enrichment was performed on log2-fold-change (log2FC) values across conditions using the positive regulation of myeloid differentiation GO term and the PAGE method described above. For differential ERV expression analysis, processed ERV annotations (see above) in FASTA format were used to build an index for Salmon workflow and then samples were processed through the Salmon-tximport-DESeq2 pipeline (see above). Upregulated ERVs were defined as p-value < 0.05 and log2FC > 2, and downregulated ERVs were defined as p-value < 0.05 and log2FC < –2. The intersections of ERV data were visualized using UpSet plots in Python^82^.

#### Reanalysis of public RNA-seq data for HL-60 derived myeloid differentiation

Ramirez et al. studied the dynamics of gene regulation in human myeloid differentiation^83^. We reanalyzed their RNA-seq data for differential gene expression changes between parental HL-60 and HL-60 derived macrophages, neutrophils and monocytes processed after 3 hours, 12 hours, 48 hours, 96 hours and 120 hours (GSE79044) using our Salmon-tximport-DESeq2 pipeline (see above). Pearson correlation coefficients were used to measure the correlation of log2-fold gene expression changes between (1) drug treatment (i.e., decitabine and RG3039 vs. DMSO) and (2) HL-60 differentiation. UpSet plots in Python^82^ were used to show specific upregulated genes in each differentiated cell type. Lastly, the onePAGE algorithm was used to assess the enrichment of select up or downregulated genes in neutrophils (see above).

#### HL-60 time-series experiments

We performed stranded RNA-seq on HL-60 cells treated with decitabine for 6 hours, 72 hours and 120 hours in duplicate (see above for experimental procedures). Differential expression analysis was performed using our Salmon-tximport-DESeq2 pipeline (see above), using a likelihood ratio test and a two-variable model design incorporating treatment condition (decitabine or DMSO) and time (6, 72 or 120 hours). Differential RNA stability analysis was performed using our STAR-featureCounts-REMBRANDTS-limma pipeline (see above). Pathway enrichment for differential expression and RNA stability data was assessed using iPAGE (see above).

#### AML cell line panel experiments

We performed stranded RNA-seq on AML cell lines treated with decitabine or DMSO for 72 hours in three replicates (see above for experimental procedures). Differential expression analysis was performed using our Salmon-tximport-DESeq2 pipeline (see above), using a Wald test. Differential RNA stability analysis was conducted using our STAR-featureCounts-REMBRANDTS-limma pipeline (see above). Pearson correlation tests from the Hmisc and corrplot R packages were used to assess correlation between differentially expressed genes in HL-60 and other AML cell lines. UpSet plots in Python^82^ were used to identify and visualize genes across multiple cell lines that conferred drug sensitivity in the CRISPRi screen (ρ score < –0.1 and p < 0.05), were RNA hypermethylated (log2FC > 1 and p < 0.05) upon decitabine treatment, and either had decreased expression or RNA stability (log2FC < –0.1 and p < 0.05) upon decitabine treatment.

### MeRIP-seq

#### Experimental procedure

We performed MeRIP-seq as previously described on HL-60 cells treated with DMSO or decitabine for 72 hours in biological duplicates^35^. First, 2 µg of the fragmented total RNA per sample was used for RNA immunoprecipitation (IP) with 5 µg of the anti-m^6^A antibody (ABE572, Millipore). RNA-seq libraries from input and IP samples were prepared using the SMARTer Pico Input Mammalian v2 RNA-seq kit (Takara) and sequenced as SE50 runs on an Illumina HiSeq4000.

#### Alignment task for mRNAs of protein coding genes and ERVs

MeRIP-seq reads were aligned to the hg38 reference genome using STAR (version 2.7.3a) with reference annotation GENCODE (version 34)^56,78^. Similarly, pre-processed annotations used to build STAR indices for each type of HERV. Then, MeRIP-seq reads were aligned separately with each STAR index to generate BAM files for the downstream tasks.

#### Experiment QC evaluations

Note that here the goal is to confirm the success of the experiment and only untreated samples are analyzed here. First, the ‘exomepeak’ function from the R package exomePeak was used to call m^6^A peaks from BAM files^84^. First, metagene plots were visualized using the Guitar R/Bioconductor package^85^. Then, the sequences of predicted m^6^A peaks were extracted using concepts described by Meng et al^84^. Briefly, the ‘bed2bed’ tool from the Computational Genomics Analysis Toolkit (with options ‘--method=merge --merge-by-namè) and the ‘getfastà module from bedtools (with options ‘-name -s -split’) were used for sequence extraction^80,81^. Finally, the FIRE algorithm was used in non-discovery mode for enrichment analysis of known m^6^A motifs (i.e., RGAC or [AG]GAC) within peak sequences, compared to randomly generated sequences^37^.

#### Peak calling and differential RNA methylation analysis

RADAR (*R*NA methyl*A*tion *D*ifferential *A*nalysis in *R*) was used to perform peak calling and differential methylation analysis^39^. Differentially methylated peaks were defined as FDR < 0.1 and logFC > 0.5. The logFC values for protein coding genes and each of ERVs used to test global hypermethylation using Wilcoxon test and t-test functions with ‘mu=0’, ‘alternative=“greater”’ options. Results are shown as annotated volcano plots using ggplot2 in R. For peak visualization across individual mRNA transcripts, the ‘plotGeneCov’ function from the RADAR R package was used to generate coverage plots. Then, the Gviz R Bioconductor package was used to draw detailed information for each mRNA transcript^86^.

Reproducible scripts for RNA methylation analyses using integrated tools are maintained as a GitHub project at https://github.com/abearab/imRIP.

### Ribo-seq

#### Experimental procedure

Ribosome profiling was performed as previously described in biological duplicates^87^. Approximately 10x10^6^ cells were lysed in ice cold polysome buffer (20 mM Tris pH 7.6, 150 mM NaCl, 5 mM MgCl2, 1 mM DTT, 100 µg/mL cycloheximide) supplemented with 1% v/v Triton X-100 and 25 U/mL Turbo DNase (Invitrogen). The lysates were triturated through a 27G needle and cleared for 10 min at 21,000 *g* at 4°C. The RNA concentrations in the lysates were determined with the Qubit RNA HS kit (Thermo). Lysate corresponding to 15 µg RNA was diluted to 200 µl in polysome buffer and digested with 0.75 µl RNaseI (Epicentre) for 45 min at room temperature. The RNaseI was then quenched by 5 µl SUPERaseIN (Thermo).

Monosomes were isolated using MicroSpin S-400 HR (Cytiva) columns, pre-equilibrated with 3 mL polysome buffer per column. 100 µl digested lysate was loaded per column (two columns were used per 200 µl sample) and centrifuged 2 min at 600 *g*. The RNA from the flow through was isolated using the RNA Clean and Concentrator-25 kit (Zymo). In parallel, total RNA from undigested lysates were isolated using the same kit.

Ribosome protected footprints (RPFs) were gel-purified from 15% TBE-Urea gels as 17-35 nt fragments. RPFs were then end-repaired using T4 PNK (New England Biosciences) and pre-adenylated barcoded linkers were ligated to the RPFs using T4 Rnl2(tr) K227Q (New England Biosciences). Unligated linkers were removed from the reaction by yeast 5’-deadenylase (New England Biosciences) and RecJ nuclease (New England Biosciences) treatment. RPFs ligated to barcoded linkers were pooled, and rRNA-depletion was performed using riboPOOLs (siTOOLs) per the manufacturer’s recommendations. Linker-ligated RPFs were reverse transcribed with ProtoScript II RT (New England Biosciences) and gel-purified from 15% TBE-Urea gels. cDNA was then circularized with CircLigase II (Epicentre) and used for library PCR. First, a small-scale library PCR was run supplemented with 1X SYBR Green and 1X ROX (Thermo) in a qPCR instrument. Then, a larger scale library PCR was run in a conventional PCR instrument, performing a number of cycles that resulted in ½ maximum signal intensity during qPCR. Library PCR was gel-purified from 8% TBE gels and sequenced on a SE50 run on an Illumina HiSeq4000.

#### Data preprocessing

The adapters in the sequencing reads were removed using cutadapt^88^ (v3.1) with options ‘--trimmed-only -m 15 -a AGATCGGAAGAGCAC’. The PCR duplicates in the reads were collapsed using CLIPflexR (v0.1.19)^89^. The UMIs for each read were extracted using UMI-tools (v1.1.1)^90^ with the options ‘extract—bc-pattern=NN’ for the 5’ end and options ‘extract --3prime --bc-pattern=NNNNN’ for the 3’ end. Reads corresponding to rRNA and other non-nuclear mRNA were removed by aligning out the reads using Bowtie2 (v2.4.2) on a depletion reference (rRNA, tRNA and mitochondrial RNA sequences)^91^. This depletion reference was built from the hg38 noncoding transcriptome (Ensembl version 96)^92,93^. The reads that did not align to the depletion reference were aligned to the hg38 mRNA transcriptome (Ensembl version 96) using Bowtie2 with options ‘-- sensitive --end-to-end --norc’. The mRNA transcriptome was built using the cDNA longest CDS reads of *Homo sapiens* downloaded from the Ensembl release version. The resulting reads were converted to BAM files and then sorted using samtools (v1.11). The duplicate reads in the sorted files were removed using UMI-tools (v1.1.1) with options ‘dedup’.

#### Differential translational efficiency (TE) analysis

Ribolog was used to compare translational efficiency across conditions (https://github.com/goodarzilab/Ribolog)^94^. Briefly, Ribolog applies a logistic regression to model individual Ribo-seq and RNA-seq reads in order to provide estimates of logTER (i.e., logFC in TE) and its associated p-value across the coding transcriptome.

### RNA expression and mutational status in cancer cell lines

RNA expression and mutational data for selected genes and cell lines were collected from the CCLE database (DepMap Public 21Q4). Cell line and gene level queries were performed using the Cancer Data Integrator^95^ – https://github.com/GilbertLabUCSF/CanDI. CanDI modified data for reproducible analysis is available at Harvard Dataverse – https://doi.org/10.7910/DVN/JIAT0H. Data were visualized in Python using the Matplotlib library.

### Multiomics data integration

To identify candidate genes among our multiomics datasets for downstream validation of our decitabine-m^6^A model, we examined the intersection of three sets of genes: (1) sensitizing hits in the CRISPRi screen, defined as ρ score < – 0.1 and p < 0.05; (2) genes with downregulated expression upon decitabine treatment, defined as log2FC < – 0.1 and p < 0.05; (3) genes with RNA hypermethylation upon decitabine treatment, defined as logFC > 1 and p < 0.05. Intersections between sets were visualized through a Venn diagram in Python.

### Quantitative RT-PCR

#### Preparation of cells for RT-qPCR and MeRIP-RT-qPCR

For each experiment, HL-60 cells were treated with DMSO or decitabine for 72 hours with three biological replicates per condition. To measure mRNA decay rates, cells were also treated with or without α-amanitin (10 µg/ml) in the final 16 hours prior to cell harvest. For MeRIP-RT-qPCR, cells were first transduced with a control sgRNA or *METTL3*-targeting sgRNA and sorted for fluorescent positive cells prior to drug treatment.

#### RNA isolation

Total RNA was isolated using the Quick-RNA Microprep kit (Zymo) with on-column DNase treatment per the manufacturer’s protocol. For MeRIP-RT-qPCR, 2 µg of the fragmented total RNA per sample was used for RNA immunoprecipitation (IP) with 5 µg of the anti-m^6^A antibody (ABE572, Millipore).

#### Quantitative RT-PCR

Transcript levels were measured using RT-qPCR by first reverse transcribing total RNA to cDNA (Maxima H Minus RT, Thermo Fisher Scientific), then using fast SYBR green master mix (Applied Biosystems) or Perfecta SYBR green supermix (QuantaBio) per the manufacturer’s instructions. HPRT1 was used as an endogenous control.

#### INTS5 primers

*Exon-junction* forward primer 5’–GGGATGTCCGCGCTGTG– 3’ and reverse primer 5’– GGACAGCTCCTGAGCACTGA–3’. *Exon-intron* forward primer 5’–GGGATGTCCGCGCTGTG– 3’ and reverse primer 5’–AGTTCTCGAGGTAGGATCCGGGT–3’. *Predicted m^6^A hypermethylated loci* forward primer 5’–TGCTGTCTGAGTTTATCCGGGCCA–3’ and reverse primer 5’– TGGACCATGCACTAATCACAGGT–3’.

#### SQLE primers

*Exon-junction* forward primer 5’–CCCAGTTCGCCCTCTTCTCGGA– 3’ and reverse primer 5’– GGTTCCTTTTCTGCGCCTCCTGG–3’. *Exon-intron* forward primer 5’– CCCAGTTCGCCCTCTTCTCGGA–3’ and reverse primer 5’–ACCTGCCGCCTTTTGCAATTCA–3’. *Predicted m^6^A hypermethylated loci* forward primer 5’–TTACTGGAGTCTGGCCGGCTCT–3’ and reverse primer 5’–CGAGTGGGTTTAAGGTTCTCCCCA–3’.

### Code availability

Reproducible code for mapping NGS reads to HERVs, flexible pathway level analysis using the PAGE algorithm, and integrated methods for MeRIP-seq analysis are publicly available at https://github.com/abearab/HERVs, https://github.com/abearab/pager and https://github.com/abearab/imRIP, respectively. Original code for all analyses in this study are available at https://github.com/GilbertLabUCSF/Decitabine-treatment.

### Miscellaneous

Subfigures and plots were generated using GraphPad Prism (GraphPad Software, La Jolla, CA), Python Matplotlib and R ggplot2. Cartoons of the dCas9 protein and sgRNA were adapted from images by the Innovative Genomics Institute, UC Berkeley and UCSF. All figures were assembled in Adobe Illustrator (Adobe, Inc.).

## SUPPLEMENTARY FIGURES

**Supplementary Figure 1.**
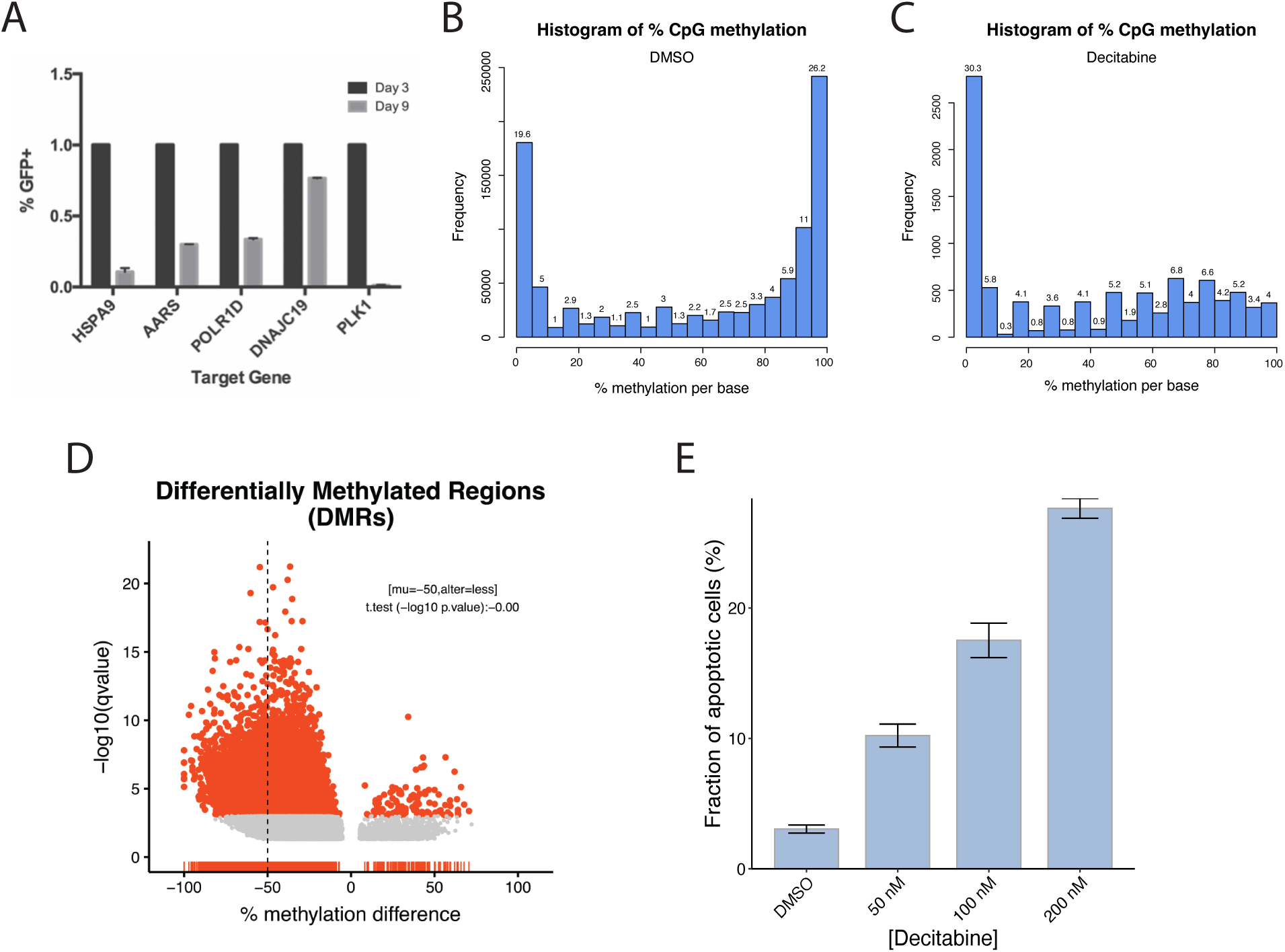
HL-60i validation and analysis of decitabine induced CpG DNA methylation changes in a public dataset. (**a**) Relative depletion of five sgRNAs targeting essential genes at day 9 (relative to day 3) in the HL-60i cell line, demonstrating functional CRISPRi activity. Each sgRNA was introduced into HL-60i via lentiviral transduction at infection rates of ∼5–20%. GFP expression was used as a surrogate for sgRNA expression and the starting infection percentage for each sgRNA was normalized to 1. Cells were monitored over time via flow cytometry. Data are shown as means ± SD for two replicates. (**b-d**) Reanalysis of a public bisulfite- sequencing dataset (GSE149954) showing frequencies of base resolution CpG methylation in HL-60 cells treated with (**b**) DMSO or (**c**) 300 nM decitabine. (**d**) Volcano plot of differentially methylated regions (DMRs) comparing cells treated with decitabine vs. DMSO. A one-sided t-test shows statistically significant global hypomethylation of DNA CpG islands. (**e**) Apoptosis assay measuring cleaved caspase 3/7 at day 5 following treatment with DMSO or decitabine. Data are shown as means ± SD for three replicates. Data were derived from the same experiment as Figure 1G.

**Supplementary Figure 2.**
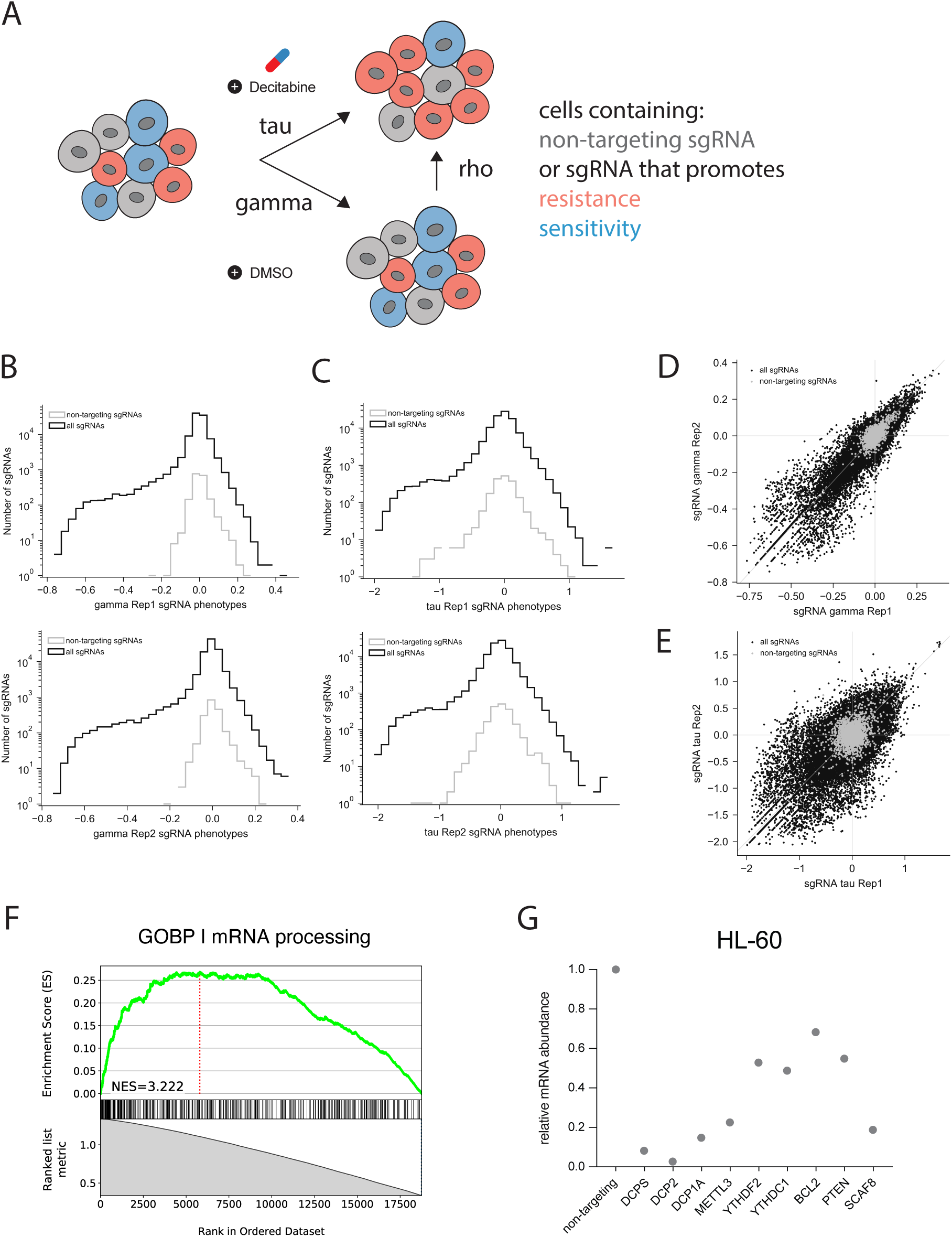
CRISPRi decitabine screen phenotype score metrics and quality control analysis for HL-60 screen. (a) Definition of CRISPRi screen phenotypes. (**b-d**) Distributions of sgRNA phenotypes per each HL-60 screen replicate show many sgRNAs are highly active relative to the negative control sgRNA distribution. (**d-e**) Scatter plots show robust correlation between HL-60 screen replicates for the gamma and tau phenotypes. Targeting and non-targeting sgRNAs included in the library are color coded black and gray, respectively. **(f)** GSEA plot showing enrichment of GO:0006397 (mRNA processing) among all screened genes ranked by Mann-Whitney p-value (corresponding to each gene’s ρ phenotype calculation). Normalized enrichment scores (NES) were calculated using the blitzGSEA Python package. (**g**) CRISPRi knockdown levels of nine hit genes in HL-60 cells. Data are plotted as mRNA abundance for each gene-targeting sgRNA relative to a non-targeting control sgRNA.

**Supplementary Figure 3.**
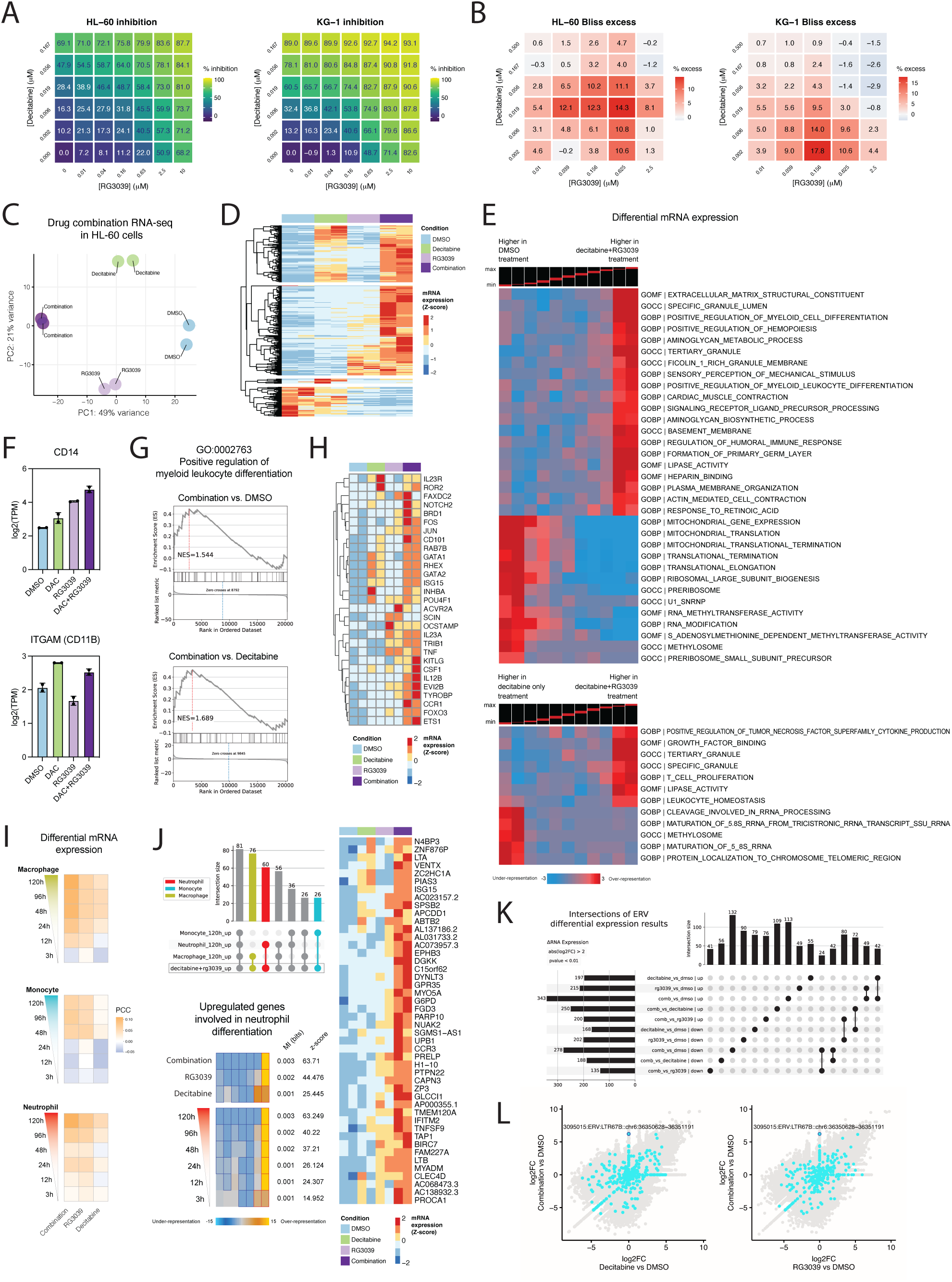
Characterizing synergy between decitabine and RG3039 in AML. (**a**) Dose response matrices for HL-60 and KG-1 treated with dose combinations of decitabine and RG3039. Heatmaps display % cell inhibition (generated using a CellTiter-Glo assay; see methods for calculations) at each dose combination. Data are shown as means of two replicates. (**b**) Bliss excess scores (i.e., observed % cell inhibition – predicted % cell inhibition assuming Bliss independence; see methods for calculations) at each dose combination. Data are shown as means of two replicates. (**c**) PCA analysis of 3’ RNA-seq (in duplicate) performed on HL-60 treated with DMSO, decitabine alone, RG3039 alone or both drugs. (**d**) DESeq2 analysis of 3’ RNA-seq data reveals differentially expressed genes. Data are shown as a heatmap displaying counts row-normalized into Z-scores. (**e**) iPAGE analysis shows enrichment of gene ontologies (GOs) (heatmap rows) among differentially expressed genes (heatmap columns) in HL-60 treated with decitabine and RG3039 (top) or decitabine alone (bottom) vs. DMSO. Genes were first ranked based on log2FC from left to right and divided into eleven equally populated bins. Red boxes show enrichment and blue boxes show depletion. For each comparison, GOs are only shown if two of the first (i.e., upregulated GO) or last (i.e., down regulated GO) bins scored above 2. (**f**) Normalized RNA-seq counts for differentiation markers *CD14* and *CD11B* in HL-60 cells treated with DMSO, decitabine, RG3039, or decitabine plus RG3039. Data are shown as means of two replicates. (**g-h**) Expression patterns for genes involved in positive regulation of myeloid leukocyte differentiation (GO:0002763). (**g**) GSEA plot shows enrichment of the GO:0002763 term in the combined drug treatment (decitabine plus RG3039) relative to DMSO or decitabine alone. Normalized enrichment scores (NES) were calculated using the blitzGSEA Python package. (**h**) Normalized counts for genes in GO:0002763 upregulated upon decitabine and RG3039 treatment. (**i**) Treatment with decitabine plus RG3039 is more highly correlated with macrophage, monocyte, and neutrophil differentiation transcriptional signatures (derived from the public dataset GSE79044) compared to treatment with either drug alone. Data are shown as correlation matrices with Pearson’s correlation coefficients (PCC). (**j**) An UpSet plots visualizes genes upregulated upon combined treatment with decitabine and RG3039 (top). PAGE analysis was performed to test for enrichment of genes involved in neutrophil differentiation, with results shown as a heatmap with rows as each logFC input and columns as cluster bins (bottom). Normalized counts for select genes most highly upregulated in the combination treatment (right). (**k**) An UpSet plot visualizes upregulated and downregulated endogenous retroviruses (ERVs) across treatment conditions. Upregulated ERVs (log2FC > 1 and p-value < 0.05) are labeled as “up”, downregulated ERVs (log2FC < −1 and p-value < 0.05) are labeled as “down” and all other ERVs are labeled as “no change”. (**l**) Scatter plots show differential ERV expression (as log2FC) in cells treated with decitabine or RG3039 alone (x-axis) vs. both drugs (y-axis). Pseudoautosomal boundary-like A (PABL_A) family members are highlighted in light blue. The labeled points correspond to the PABL_A chr9:9641512-9642657 locus, which is only upregulated in the decitabine and RG3039 drug combination.

**Supplementary Figure 4.**
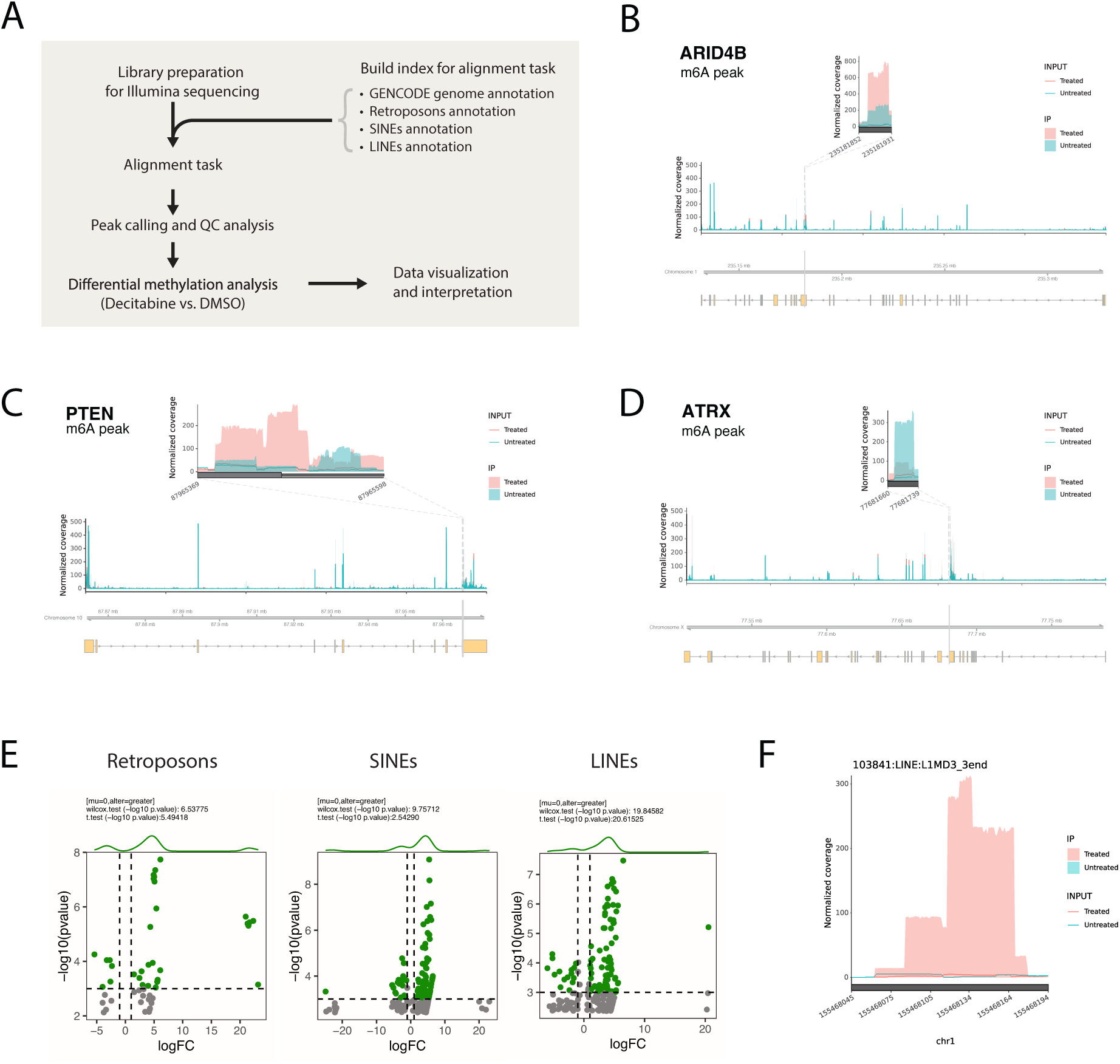
MeRIP-seq workflow to identify differentially methylated peaks associated with decitabine treatment in HL-60 cells. (**a**) Schematic of MeRIP-seq computational workflow. (**b-d**) Visualization of m^6^A peaks across mRNA transcripts of **(b)** *ARID4B*, **(c)** *PTEN* and **(d)** *ATRX*. Peaks were called using the RADAR algorithm and plots were generated using the RADAR and Gviz R packages. MeRIP-seq experiments were performed in biological duplicates for each condition. **(e)** Differential methylation analysis shows significant changes in RNA methylation peaks in HL-60 cells treated with decitabine relative to DMSO. Global hypermethylation is observed in the decitabine condition for different families of ERVs. Peaks are called using the RADAR algorithm and visualized as annotated volcano plots. Wilcoxon and t-tests are used to assess statistical significance of global hypermethylation. **(f)** Coverage plot for a representative hypermethylated peak in the L1MD3_3end LINE transcript upon decitabine treatment (pink) compared to DMSO control (blue).

**Supplementary Figure 5.**
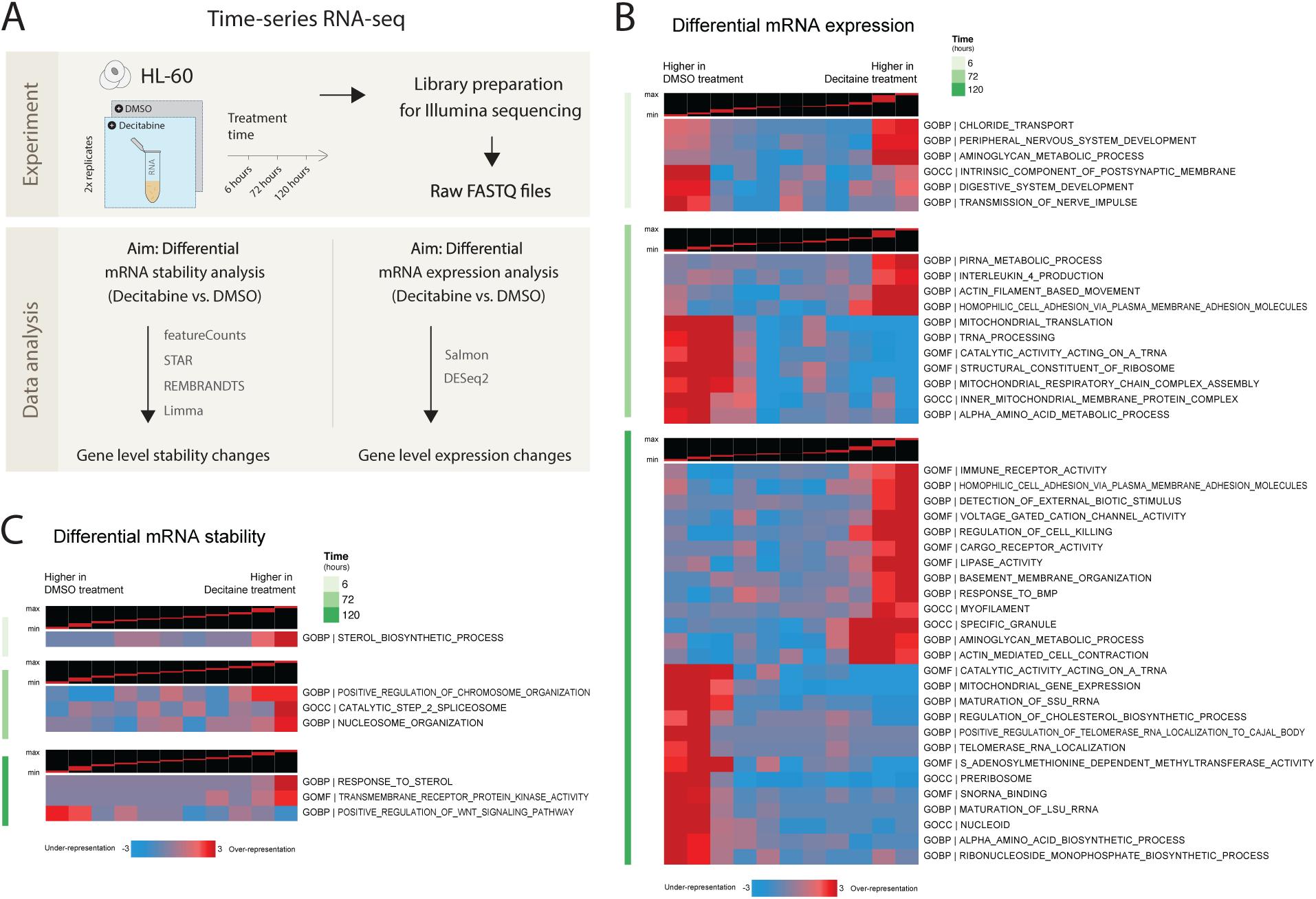
Pathway-level changes in mRNA expression and stability associated with decitabine treatment in HL-60 cells. (**a**) Schematic of RNA-seq workflows in HL-60 cells. Two parallel workflows describe analysis of differential mRNA stability (left) and differential mRNA expression (right). (**b-c**) Gene set enrichment analysis with the iPAGE algorithm shows enrichment of GOs (heatmap rows) among changes in (**b**) RNA stability and (**c**) gene expression (heatmap columns; ranked and quantized into equal bins) upon decitabine treatment. The logFC values for HL-60 cells treated with decitabine vs. DMSO at 6 hours (top), 72 hours (middle) and 120 hours (bottom) were assessed separately. Highly-enriched GOs with genes upregulated or downregulated upon decitabine treatment are shown.

**Supplementary Figure 6.**
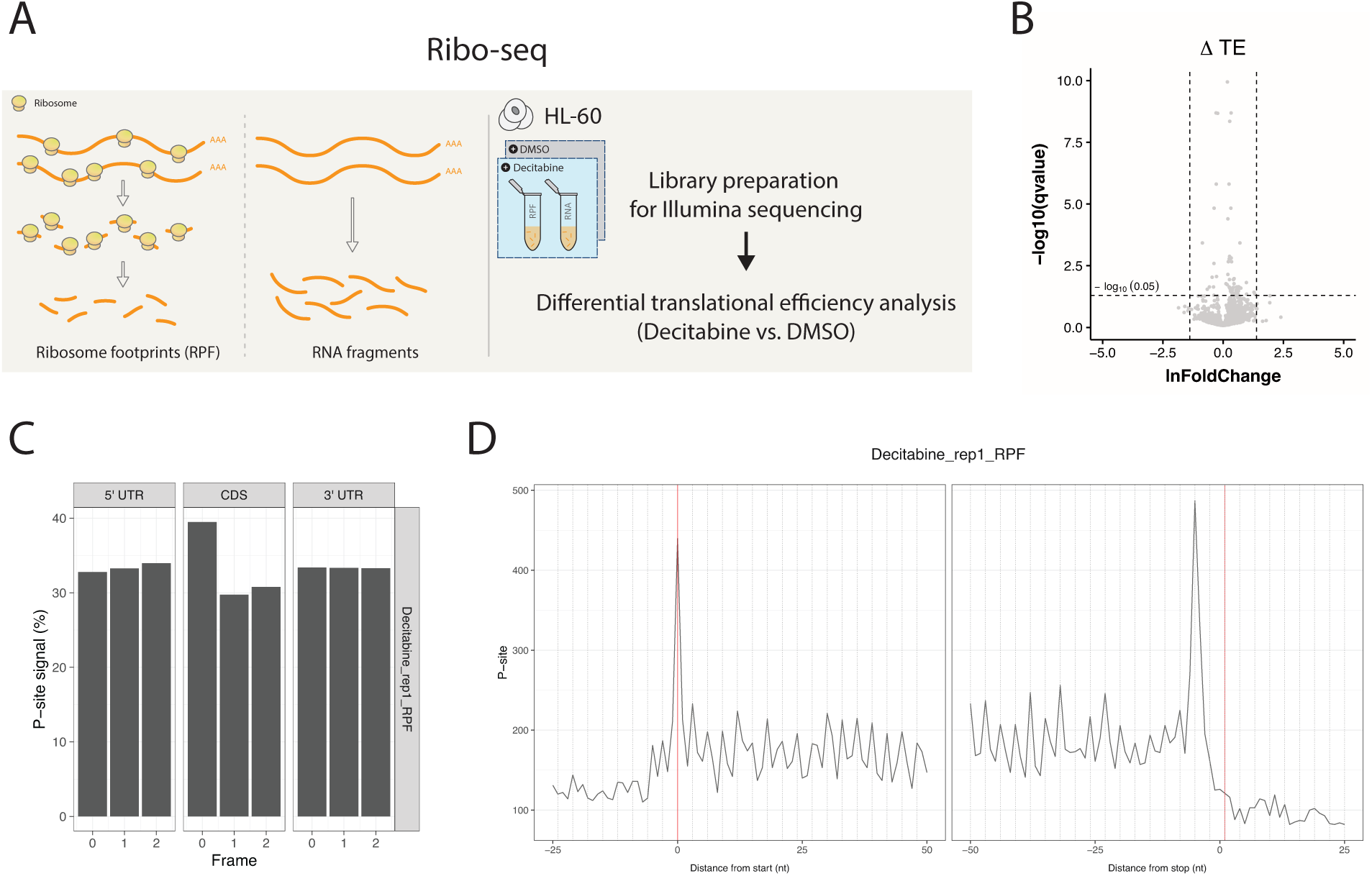
Translational efficiency (TE) changes associated with decitabine treatment in HL-60 cells. (**a**) Schematic of Ribo-seq experimental workflow. (**b**) Volcano plot visualization of Ribolog-calculated translational efficiency ratios (TERs) between the decitabine and DMSO conditions. (**c**) Bar plots showing enrichment of P-sites in the first frame of coding sequence (CDS) but not UTRs, consistent with ribosome protected fragments derived from protein coding mRNAs. (**d**) Ribosome occupancy profiles based on the 5’ and 3’ reads mapped to a reference codon for one sample (decitabine treated HL-60, single replicate).

**Supplementary Figure 7.**
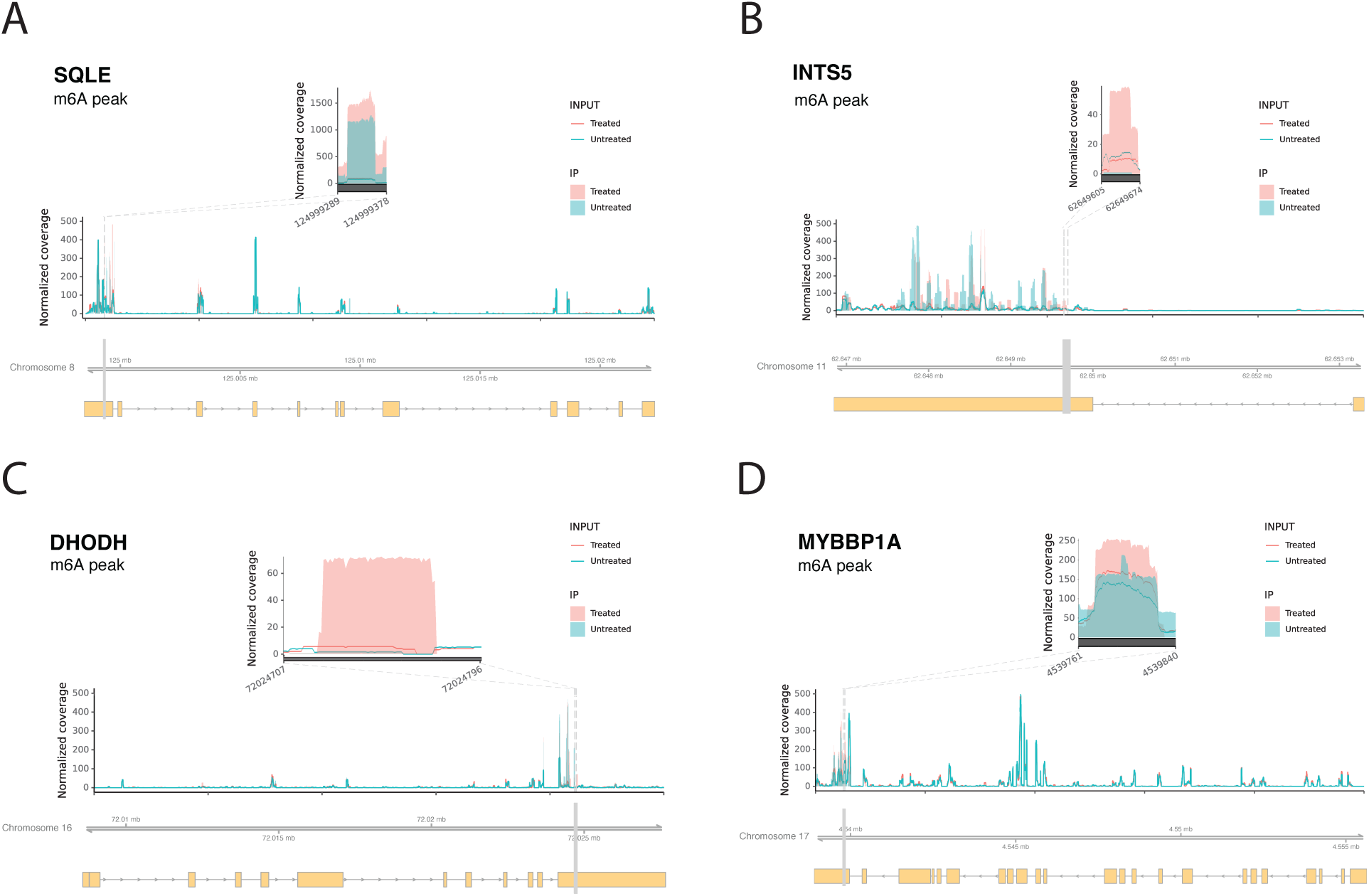
RNA m^6^A hypermethylated peaks from MeRIP-seq in HL-60 following decitabine treatment. (**a-d**) Visualization of m^6^A peaks across mRNA transcripts of **(a)** *SQLE*, **(b)** *INTS5*, **(c)** *DHODH* and **(d)** *MYBBP1A*. Peaks were called using the RADAR algorithm and plots were generated using the RADAR and Gviz R packages. MeRIP-seq experiments were performed in biological duplicates for each condition.

**Supplementary Figure 8.**
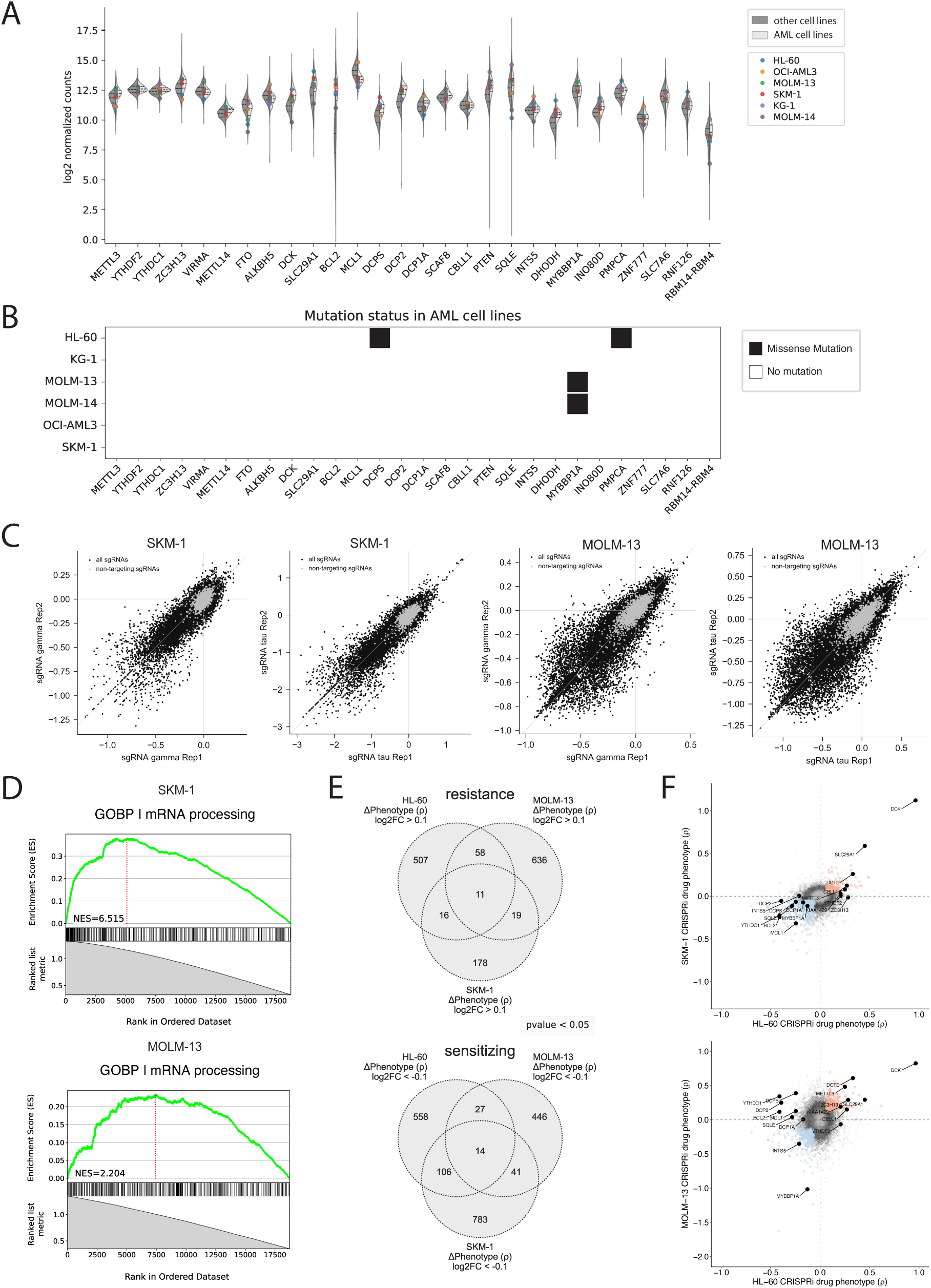
Analysis of SKM-1 and MOLM-13 cell lines and genome-scale CRISPRi decitabine screens: quality control and comparisons to the HL-60 screen. (**a**) RNA expression levels for genes of interest (shown as log2 normalized counts) across AML cell lines vs. other cancer types using the CCLE database (DepMap Public 21Q4) curated with Cancer Data Integrator (CanDI). In total, 54 AML cell lines and 1,771 other cancer type cell lines are shown, with the 6 AML cell lines used in this study highlighted. (**b**) Mutational status of genes of interest across the 6 AML cell lines used in this study. (**c**) Scatter plots show robust correlation between replicates for the gamma and tau phenotypes in SKM-1 (left) and MOLM-13 (right) genome-scale CRISPRi decitabine screens. (**d**) GSEA plots for the SKM-1 (top) and MOLM-13 (bottom) screens show enrichment of the GO:0006397 (mRNA processing) term among all screened genes ranked by Mann-Whitney p-value (corresponding to each gene’s ρ phenotype calculation). Normalized enrichment scores (NES) were calculated using the blitzGSEA Python package. (**e**) Venn diagrams of significant hits across screens in three AML cell lines show overlapping and cell-line specific resistance (top) and sensitizing (bottom) phenotypes. Hits were selected by absolute gene-level rho (ρ) score values above 0.1 and Mann-Whitney p-values less than 0.05. (**f**) Scatter plots of gene-level rho (ρ) scores comparing the HL-60 screen to the SKM-1 (top) and MOLM-13 (bottom) screens. Several hits of interest shared across cell lines are labeled in black.

**Supplementary Table 1. HL-60 CRISPRi decitabine screen**

Gene-level phenotype scores and sgRNA protospacer sequences for validation assays.

**Supplementary Table 2. Pathway-level analysis of HL-60 CRISPRi drug phenotype scores**

Gene set enrichment analysis (GSEA) results using gene ontology (GO) gene sets. Two distinct GSEA analyses were performed (see methods).

**Supplementary Table 3. Differential RNA methylation analysis**

Differential analysis of MeRIP-seq data with RADAR (decitabine vs. DMSO).

**Supplementary Table 4. SKM-1 and MOLM-13 CRISPRi decitabine screens**

Gene-level phenotype scores for each screen and comparison of CRISPRi drug phenotype across three AML cell lines.

**Supplementary Table 5. Pathway-level analysis of AML cell lines CRISPRi drug phenotype scores**

Merged results from gene set enrichment analysis (GSEA) using gene ontology (GO) gene sets across three AML cell lines. Two distinct GSEA analyses were performed (see methods).

